# RNA-dependent RNA polymerase of predominant human norovirus forms liquid-liquid phase condensates as viral replication factories

**DOI:** 10.1101/2023.08.24.554692

**Authors:** Soni Kaundal, Ramakrishnan Anish, B. Vijayalakshmi Ayyar, Sreejesh Shanker, Gundeep Kaur, Sue E. Crawford, Jeroen Pollet, Fabio Stossi, Mary K. Estes, B.V. Venkataram Prasad

## Abstract

Many viral proteins form biomolecular condensates via liquid-liquid phase separation (LLPS) to support viral replication and evade host antiviral responses, and thus, they are potential targets for designing antivirals. In the case of non-enveloped positive-sense RNA viruses, forming such condensates for viral replication is unclear and less understood. Human noroviruses (HuNoV) are positive-sense RNA viruses that cause epidemic and sporadic gastroenteritis worldwide. Here, we show that the RNA-dependent-RNA polymerase (RdRp) of pandemic GII.4 HuNoV forms distinct condensates that exhibit all the signature properties of LLPS with sustained polymerase activity and the capability of recruiting components essential for viral replication. We show that such condensates are formed in HuNoV-infected human intestinal enteroid cultures and are the sites for genome replication. Our studies demonstrate the formation of phase separated condensates as replication factories in a positive-sense RNA virus, which plausibly is an effective mechanism to dynamically isolate RdRp replicating the genomic RNA from interfering with the ribosomal translation of the same RNA.

**Teaser:** Polymerase of a positive-sense RNA virus forms LLPS to regulate replication as an elegant solution for an enigmatic question.

## Introduction

The compartmentalization of biomolecules is essential for efficiently regulating numerous biological processes in a dense cellular environment. A wealth of new research has shown that, apart from the traditional membrane-bound organelles, eukaryotic cells also contain membraneless organelles such as P granules, nucleoli, Cajal bodies, etc. (*1–3*). These membraneless organelles, also known as biomolecular condensates, are formed by the process of liquid-liquid phase separation (LLPS), where a single, uniform liquid phase separates into a dense phase that coexists with the surrounding dilute phase (*4, 5*). These condensates, often initiated by a single protein, selectively allow the incorporation of proteins and nucleic acids to provide exclusive compartments for performing specialized cellular functions, including gene regulation, protein synthesis, and signaling (*5, 6*). Depending upon the sequence and concentration of the protein, the presence of other biomolecules, and the physiochemical properties of the surroundings, the dense phase can have the material properties of a liquid, gel, or solid (*3, 7, 8*). The liquid-like condensates typically have a spherical shape with a dynamic interior and exhibit the ability to fuse with each other, deform upon physical contact, and revert to the spherical shape (*3, 8*). Signature features of the proteins undergoing LLPS include the presence of a nucleic acid binding domain, a disordered/flexible region, a low complexity region, and the ability to form oligomers (*9, 10*).

Many viruses, upon infecting host cells, form distinct intracellular compartments, often referred to as viroplasms or replication factories (RFs), for genome replication and capsid assembly (*11, 12*). In negative-sense RNA and dsRNA viruses, these compartments are typically membraneless organelles within the cytoplasm, commonly known as inclusion bodies. A growing amount of evidence shows that these inclusion bodies have liquid-like properties. The formation of liquid condensates by viral proteins (N and P) was initially demonstrated for Mononegavirales with negative-strand RNA genome, including rabies virus (*13*), vesicular stomatitis virus (*14*), measles virus (*15*), and respiratory syncytial virus (*16*). In the case of rotavirus, a dsRNA virus, non-structural protein NSP5, in association with the NSP2 protein, undergoes LLPS to form viroplasms, the sites for viral replication(*17*). In contrast, the viral replication complexes of positive-sense RNA viruses are associated with modified cellular membranes (*18, 19*). However, recent research has demonstrated that proteins encoded by positive-sense RNA viruses can also exhibit phase separation behavior and play a critical role in the viral life cycle. For example, phase separation by the N protein of SARS-CoV2 is implicated in virus replication, genome packaging, and immune suppression (*20–25*). This implies that other positive-sense RNA viruses likely depend on crucial proteins for phase separation to control replication, and discovering these proteins could pave the way for improved drug therapies.

Apart from the N protein of SARS-CoV2, there are very limited reports of proteins encoded by other positive-sense RNA viruses that undergo phase separation (*26*), particularly in the context of their replication. Here, we investigated the phase separation behavior of protein (s) encoded by human norovirus (HuNoV), a positive-sense RNA virus. HuNoVs are the leading cause of viral gastroenteritis and pose a significant global health and economic burden without any approved vaccines or antivirals (*27, 28*). These icosahedral, single-stranded RNA viruses, belonging to the *Norovirus* genus in the *Caliciviridae* family account for an estimated ∼685 million cases and ∼212,000 deaths globally per year, with an economic impact of $60 billion (*27–31*). The NoVs are classified into at least 10 genogroups (GI-GX) and 49 genotypes (*32*). NoVs in genogroups GI, GII, GIV, GVIII, and GIX infect humans (*32, 33*), and among these, HuNoVs belonging to genogroup II and genotype 4 (GII.4) are responsible for ∼80 % of the outbreaks worldwide (*34*). The NoV genome is covalently linked to VPg and consists of three open reading frames (ORFs). ORF1 encodes a polyprotein that is cleaved into six nonstructural proteins by the viral protease to produce p48 (NS1-2), p41 (NS3), p22 (NS4), VPg (NS5), protease (NS6), and the RNA-dependent RNA polymerase (RdRp) (NS7) (*35*). ORF2 and ORF3 encode the major capsid protein VP1 and a minor structural protein VP2, respectively (*^36^*). Our understanding of the HuNoV replication processes in infected cells has been hampered by the lack of a robust cell culture system. Much of our current understanding of the entry and replication processes of HuNoVs has come from using the human intestinal stem cell-derived, nontransformed enteroid (HIE) monolayer cultures (*^37–39^*) for virus cultivation.

Using bioinformatics analysis, we found that among the nonstructural proteins encoded by the pandemic GII.4 Sydney strain, RdRp has the highest propensity to undergo LLPS to form biomolecular condensates. In addition to the biophysical properties of GII.4 RdRp, such as its modular architecture, the presence of an N-terminal disordered/flexible region, the ability to homo-oligomerize and bind RNA, and the critical role it plays during viral replication prompted us to experimentally test the hypothesis that GII.4 RdRp drives the formation of biomolecular condensates that are conducive for viral replication through LLPS. Our studies show that RdRp, consistent with its biophysical properties, indeed forms highly dynamic liquid-like condensates via LLPS at physiologically relevant conditions and that the disordered/flexible N-terminal region of RdRp is critical for LLPS. Our studies further show that distinct puncta observed in the virus-infected HIEs are liquid-like condensates representing replication compartments, as evidenced by the presence of RdRp, replication intermediate dsRNA, and nonstructural protein (VPg). Together, these findings establish that HuNoV RdRp undergoes LLPS to form biomolecular condensates, which serve as efficient platforms for viral replication.

## Results

### HuNoV RdRp exhibits the obligatory properties to undergo LLPS

We used the DeePhase server (*40*) to analyze the primary amino acid sequences of all the nonstructural proteins encoded by HuNoV GII.4 (Sydney) for their ability to undergo LLPS (Fig. 1A). This analysis predicted that RdRp, with an overall score of 0.67, has the highest propensity to undergo LLPS. The known biochemical and enzymatic properties of RdRp are consistent with this prediction. As a protein critical for the transcription and replication of genomic RNA, the RNA binding property of RdRp satisfies one of the criteria for undergoing LLPS. Other requirements include the presence of disordered or flexible regions and the ability to form higher-order oligomers. Analysis by PONDR (Fig. 1B) (*41*) and IUPred2 (Fig. 1C) (*42*) software shows N-terminal ∼50 residues in RdRp are disordered.

**Figure 1.**
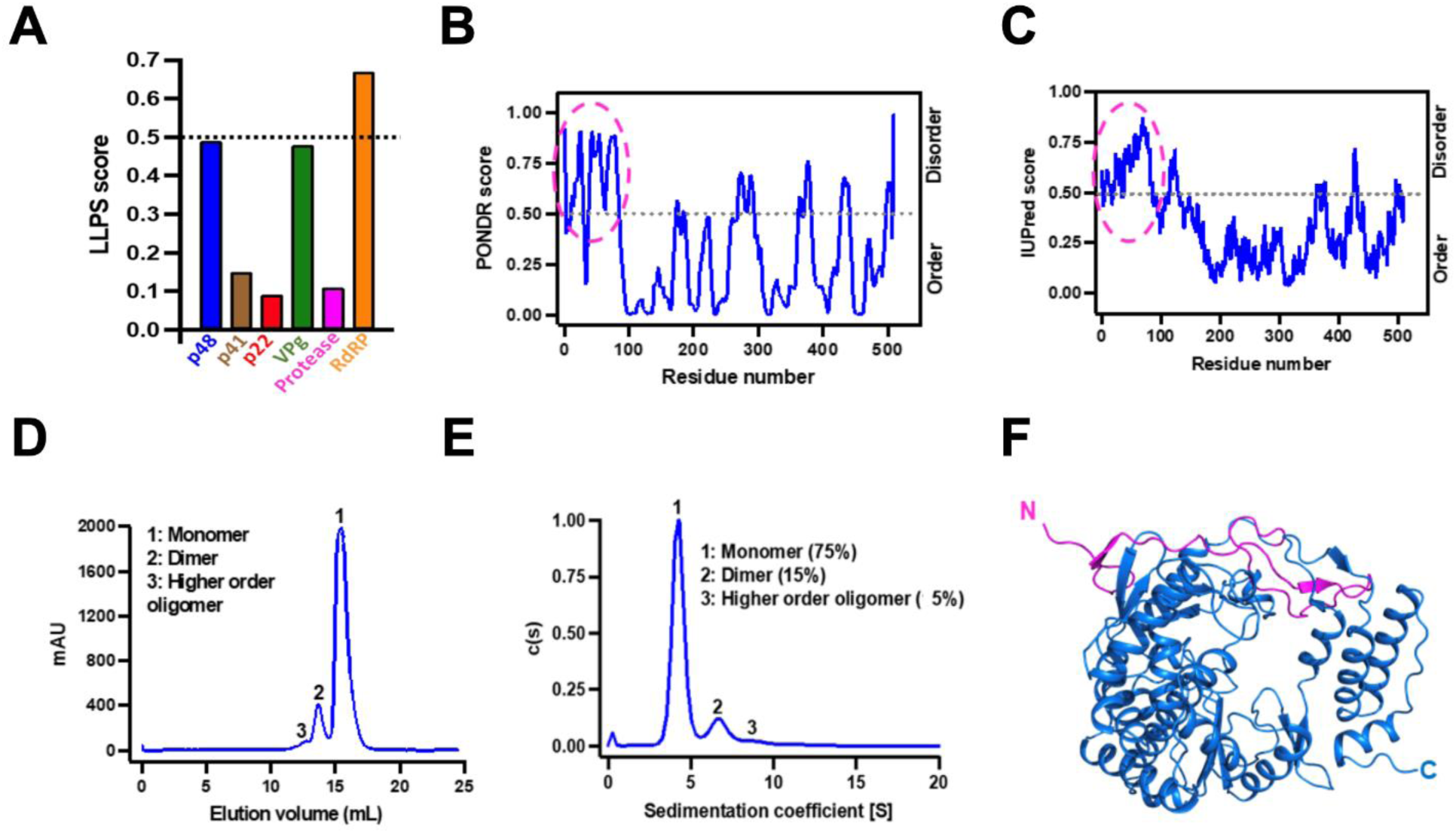
GII.4 RdRp has the required properties to undergo LLPS. (A) Bioinformatics analysis of the primary amino acid sequences of all nonstructural proteins of HuNoV GII.4 to predict LLPS propensity using DeePhase. The dotted black line indicates the threshold LLPS score of proteins to undergo LLPS. (B-C) The disorder prediction of GII.4 RdRp primary amino acid sequence using bioinformatics tools (B) PONDR (C) IUPred2. The dotted black line in both panels indicates the threshold disorder score and the dotted pink circle shows the predicted disordered N-terminal region. (D-E) The oligomeric state of GII.4 RdRp analyzed using (D) size exclusion chromatography and (E) sedimentation velocity analytical ultracentrifugation. (F) A cartoon representation of the crystal structure of GII.4 RdRp showing the disordered/flexible N-terminal region colored in pink.

To experimentally confirm the LLPS driving properties of GII.4 RdRp, we purified recombinant RdRp and determined its oligomeric state using size exclusion chromatography (SEC) (Fig. 1D) and sedimentation velocity analytical ultracentrifugation (SV-AUC) (Fig. 1E) which show that while most of the RdRp (75 %) exists as a monomer in physiologically relevant conditions, a fraction of it also tends to form dimers (15 %) and higher-order oligomers. Both monomers and the higher-order oligomers (dimers) exhibited comparable polymerase activity (Fig. S1). We also determined the X-ray crystal structure of RdRp at 2.0 Å resolution using molecular replacement (PDB ID: 8TUF, Supplementary Table 1). The crystal structure shows, as expected, all the features observed in the previous crystallographic structures of the NoV RdRps (*43–46*), with a polypeptide fold resembling a partially closed right hand, with fingers, thumb, and palm as subdomains (Fig. 1F). The N-terminal 50 residues with only two small stretches of β-stands form a flap connecting the finger and the thumb domains. To visualize the possible flexible regions in the RdRp, we performed Size Exclusion Chromatography coupled to Small Angle X-ray Scattering (SEC-SAXS) studies combined with molecular dynamics simulations using BILBOMD (*47*). Guinier plot analysis suggests that purified RdRp does not show any signs of aggregation and has a radius of gyration (R_g_) ∼25 Å (Fig. S2A and B). Kratky plot shows that RdRp is globular in shape and exists as a monomer in solution (Fig. S2C), and the P(r) plot shows that RdRp is ∼70 Å in diameter (D_max_) (Fig. S2D). To model the population of different conformations adopted by RdRp in solution, we used the X-ray structure of RdRp (PDB ID: 8TUF) as the template and generated an ensemble of molecular models (∼10,000) varying in R_g_ and D_max_ (Fig. S2E) and the topmost models with χ^2^ close to 1 were further used for calculating theoretical SAXS curves. The experimental scattering profile obtained from purified RdRp was compared to the theoretical scattering profiles of top 8 BILBOMD atomistic models (Fig. S2E). We superposed the shortlisted BILBOMD models to the experimentally determined crystal structure of GII.4 RdRp and observed that fingers, thumb, and palm domains of BILBOMD models superpose well (rmsd: 1.05 Å over 465 Cα atoms) except the N-terminal 50 residues, which appear to adopt a wide range of orientations in the solution (Fig. S2F) indicating its flexible nature. Together, these studies suggest that RdRp is a suitable candidate to further investigate its phase separation behavior.

### HuNoV RdRp undergoes phase separation to form dynamic liquid-like condensates

To experimentally validate that RdRp undergoes LLPS, we studied its phase separation behavior in vitro using light scattering experiments and confocal microscopy. Under physiological conditions, the homogeneously mixed purified RdRp (10 μM) phase separates to form a demixed turbid solution when incubated at 37 °C for 45 min (Fig. S3A). The presence of condensates in the phase separated solution was examined by confocal microscopy using a mixture of Atto 488-labeled (1 %) and unlabeled (99 %) RdRp using depression slides. The micrographs showed the formation of distinct spherical condensates (Fig. 2A). We examined the liquid-like nature of these condensates by assessing their sensitivity towards the aliphatic alcohol 1,6-hexanediol (1,6 HD), which is commonly used as a chemical probe to distinguish between liquid-like and gel-like states of biomolecular condensates (*48, 49*). The 1,6 HD (5%) treatment shows the dissolution of condensates, indicating their liquid-like nature (Fig. 2B). Further, we investigated the dynamic nature of these condensates using fluorescence recovery after photobleaching (FRAP) of the RdRp molecules in the condensates. We observed a complete fluorescence recovery (98 %) with a half-life (t_1/2_) of 53.3 ± 2.1 s (Fig. 2C, D and Supplementary Video 1) indicating diffusion of RdRp molecules in the condensate. Using time-lapse confocal microscopy, we observed that smaller RdRp condensates fuse to form a larger condensate (Fig. 2E and Supplementary Video 2) and undergo surface wetting and dripping (Fig. 2F). They further undergo dissolution upon dilution, indicating their reversible nature (Fig. S3B). Our studies thus clearly show that HuNoV GII.4 RdRp undergoes phase separation to form dynamic, liquid-like condensates, which are dense and high in RdRp concentration yet allow the RdRp molecules to move freely within the condensates and exchange with the surrounding environment.

**Figure 2.**
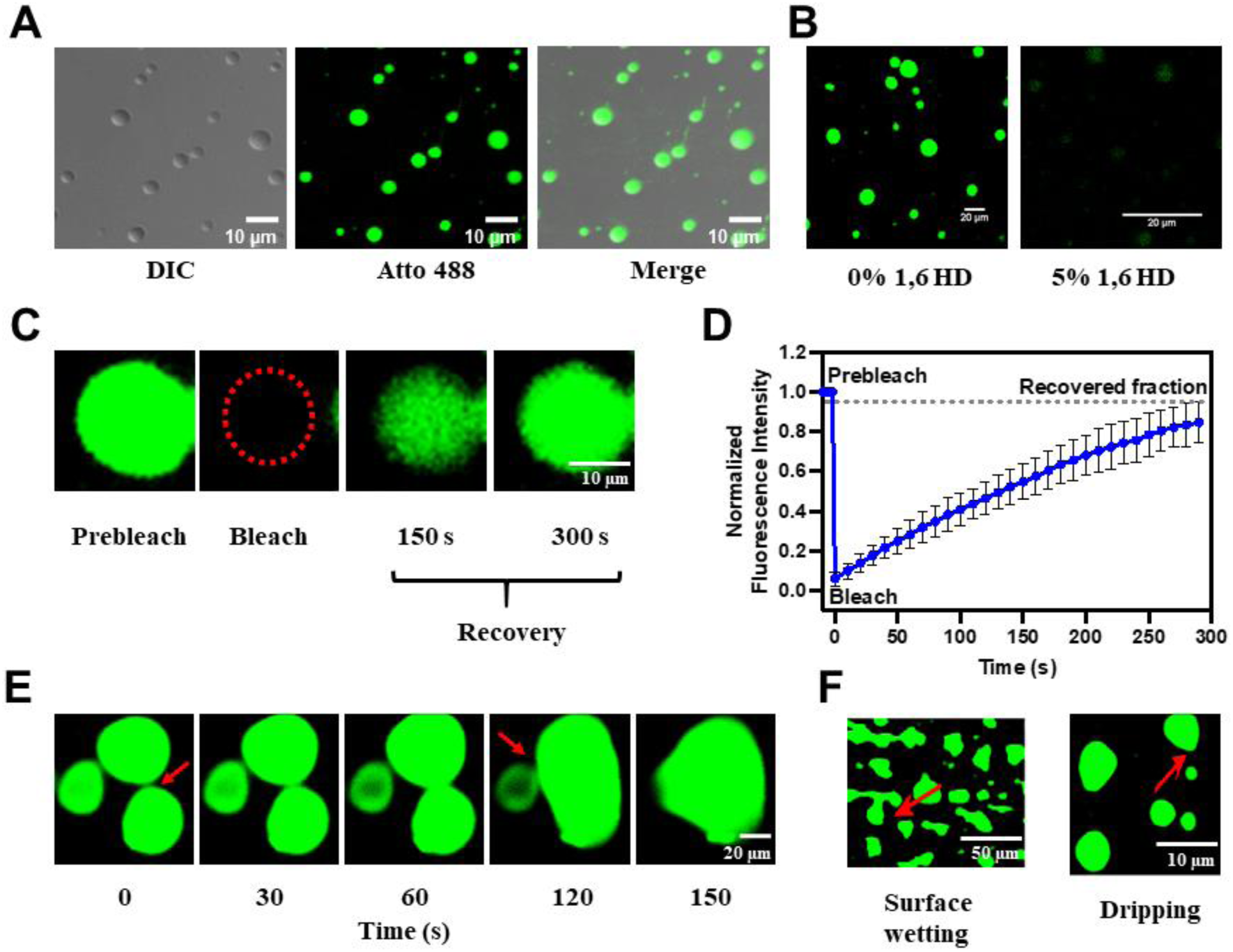
HuNoV GII.4 RdRp undergoes LLPS in vitro. (A) Differential interference contrast (DIC) and confocal images of phase separated RdRp (1 % Atto 488 labeled and 99 % unlabeled). Phase separated RdRp possesses liquid-like properties. (B) Effect of 1,6 HD on LLPS of RdRp. (C) FRAP of RdRp in condensate photobleached at the position indicated by the red circle. (D) The graph shows the recovery curve where normalized fluorescence intensity was plotted against time. (E) Time-lapse images of RdRp condensates exhibit the formation of larger condensates by the fusion of smaller condensates over time. The red arrows indicate the site of fusion. (F) Surface wetting and dripping using confocal microscopy. The red arrows indicate the position of surface wetting and dripping.

### Molecular crowding, pH, and salt concentration modulate phase separation properties of HuNoV GII.4 RdRp

To examine whether there is a minimum concentration of RdRp necessary to undergo LLPS, and how this concentration is affected by molecular crowding agents such as polyethylene glycol (PEG), typically used to simulate intracellular molecular crowding, we used two different techniques: light scattering experiments to measure turbidity (Fig. 3A) and confocal microscopy to assess LLPS formation (Fig. 3D). Different concentrations (0.5-25 μM) of RdRp were screened with and without PEG3350 (5-20 %). Confocal microscopy results suggest the minimum concentration for phase separation of RdRp decreases from 5 μM to 0.5 μM by adding 10 % PEG3350, consistent with the findings from our light scattering experiments (Fig. 3A). Similar results were obtained with higher molecular weight 5% PEG8000 (Fig. S4). Together, these observations indicate that an increase in molecular crowding, as expected inside the cells during virus replication, decreases the minimum concentration of the protein to phase separate.

**Figure 3.**
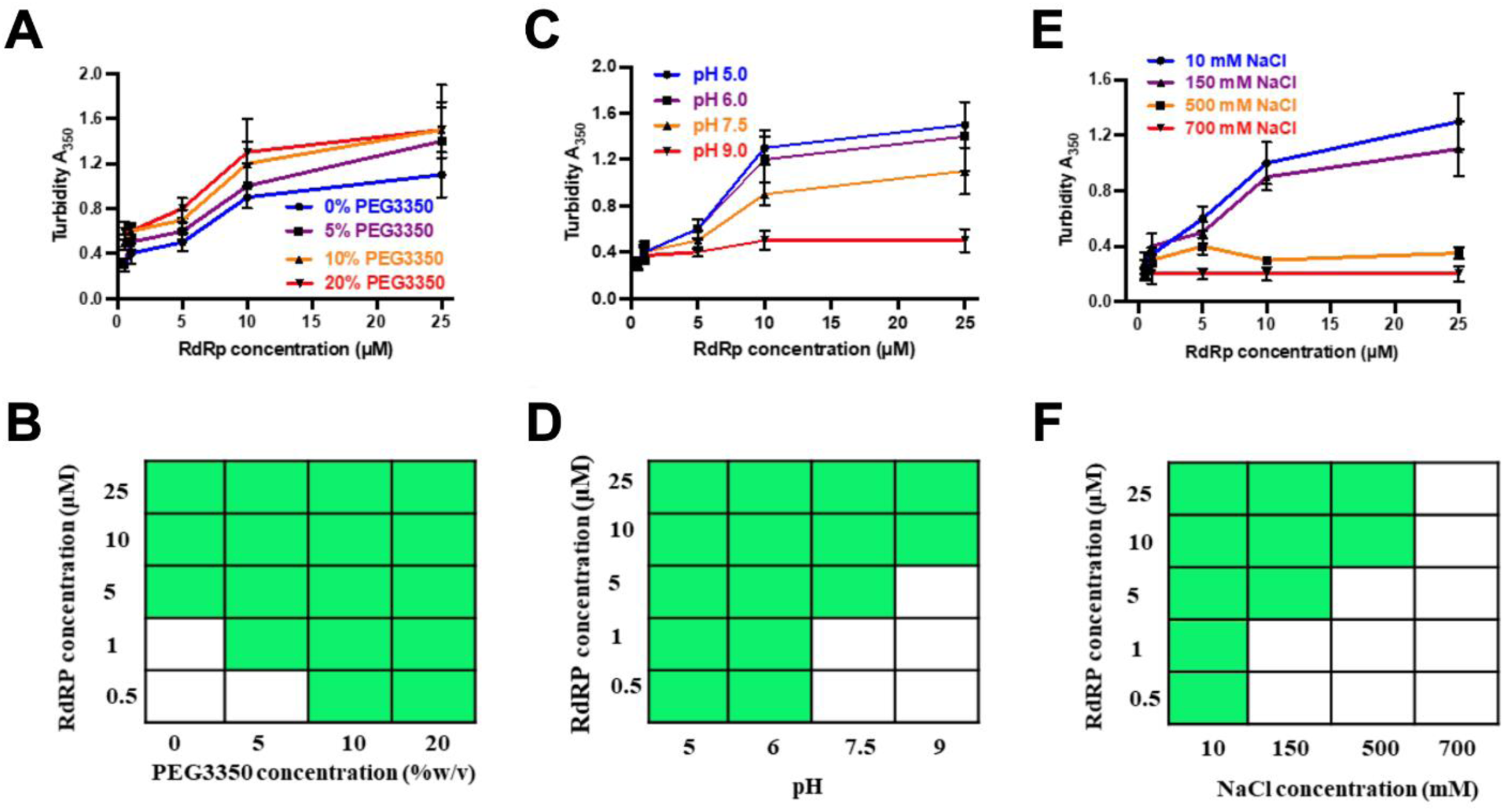
Phase separation of GII.4 RdRp is modulated by PEG3350 concentration, pH, and salt concentration. Effect of (A) protein and PEG3350 concentrations, (B) protein concentrations and pH, (C) protein and salt concentrations on the phase separation of RdRp was assessed by measuring the turbidity of the samples at OD 350 nm. (D-F) A phase diagram of RdRp at (D) protein and PEG3350 concentrations, (E) at protein concentrations and pH, at (F) protein and salt concentrations constructed from the confocal microscopy images. The green boxes indicate LLPS (condensate state), and the white boxes indicate no LLPS (soluble state).

As it has been shown that virus infection can alter the intracellular pH and salt concentrations of the host cell (*50–52*), we further examined if these factors can affect the phase separation behavior of RdRp. We observed increased turbidity in the purified RdRp at an acidic pH of 5.5 compared to a neutral pH, indicating that an acidic pH promotes phase separation more effectively than a neutral pH. In contrast, the turbidity was notably less at an alkaline pH of 9.0 (Fig. 3B). Since turbidity measurement is affected by the condensate’s size, shape, and density, we further performed confocal microscopy at different pH conditions (Fig. 3E). RdRp forms larger condensates at pH 5.5 than at pH 7.5, and, at pH 9, the condensates are few and have an irregular shape. In addition, at pH 5.5, the minimum concentration of RdRp required to form LLPS decreased to 0.5 μM from 5 μM at pH 7.5. To understand the conformational behavior of RdRp at different pH, we performed the AUC. The results show that at pH 7.5, 75% of RdRp is a monomeric and the rest is dimers and higher-order oligomers. At pH 9.0, RdRp loses its ability to form dimers and higher-order oligomers. Interestingly, at pH 5.5, most of the RdRp forms dimers and higher-order oligomers (Fig. S5). These results suggest that pH plays an important role in the oligomerization of RdRp and possibly explains why a lesser concentration of RdRp is required for condensate formation at pH 5.5 compared to pH 7.5.

We also tested the effect of ionic strength on the phase separation of RdRp by measuring the turbidity of the solutions containing varying concentrations of NaCl using turbidity measurements at 350 nm (Fig. 3C) and by imaging the condensates using confocal microscopy (Fig. 3F). These studies showed GII.4 RdRp undergoes phase separation at NaCl concentrations between 10-500 mM. Recent studies have shown a similar behavior of the N protein of SARS-CoV-2 concerning pH and ionic strength (*23*).

### The flexible N-terminal region is critical for HuNoV GII.4 RdRp LLPS

To validate the predicted role of the flexible N-terminal region of RdRp in forming the condensates, first, we analyze the N-terminal deletion protein, HuNoV GII.4 RdRp-NΔ51 using DeePhase (*40*). This analysis shows that deleting N-terminal 51 residues reduces the DeePhase score to 0.52 (Fig. S6A). Analysis of RdRp using ParSe (*53*), another bioinformatics tool to predict the amino acid involved in LLPS, suggests that the N-terminal region of RdRp plays an important role in LLPS (Fig. S6B). We expressed and purified RdRp-NΔ51 and investigated if this protein is properly folded and enzymatically active. The CD spectra of RdRp wild type (WT) and RdRp-NΔ51 showed two distinct negative maxima at λ = 208 nm and 220 nm and a strong positive peak around 190 nm indicative of an α-helical conformation and thus of proper folding of the N-terminal deletion construct (Fig. 4A). We also assessed the RNA polymerization activity of RdRp-NΔ51 protein with a fluorescence-based assay using SYTO9 dye used in many recent studies (*54, 55*). The results indicate that RdRp-NΔ51 is enzymatically active with comparable activity with RdRp WT (Fig. 4B and C, respectively). Size exclusion chromatography of this deletion construct shows a reduction in their ability to form oligomers (Fig. 4D). Confocal microscopy images show that deletion of N-terminal 51 residues (RdRp-NΔ51), showed few and very small condensates that were difficult to image compared to RdRp WT (Fig. 4E). Together these results demonstrate that flexible N-terminal region of RdRp plays a critical role in its phase separation behavior.

**Figure 4.**
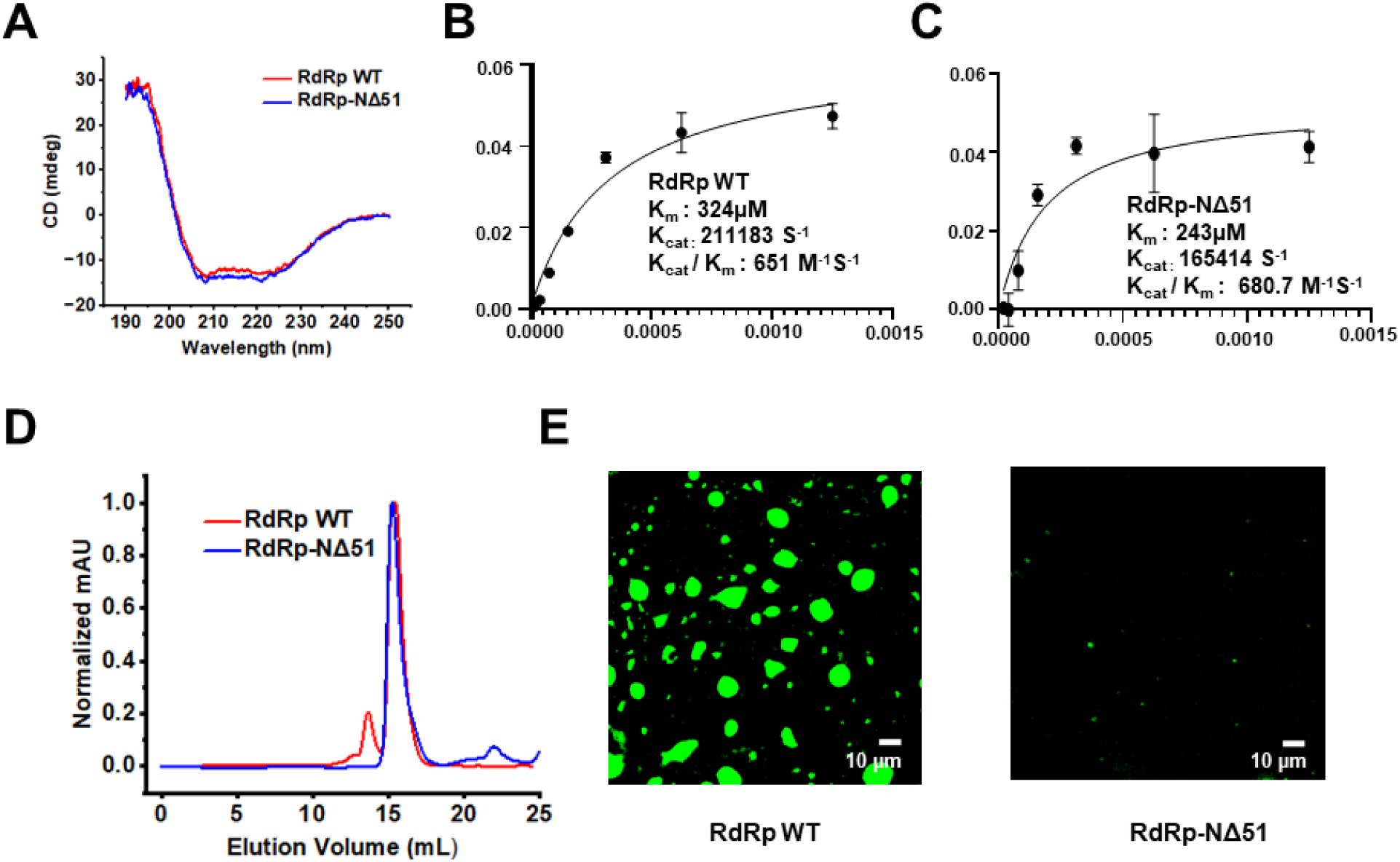
Effect of N-terminal deletion on phase separation of GII.4 RdRp. Purified RdRp WT and RdRp-NΔ51 were analyzed by **(A)** CD spectroscopy **(B and C)** Enzyme activity **(D)** Size exclusion chromatography and **(E)** confocal microscopy.

### RdRp is enzymatically active within liquid condensates

We estimated the concentration of RdRp in the condensates using the centrifugation method (*56*), which shows that it is ∼20 times higher than the surrounding dilute phase. To examine if the highly concentrated protein in the condensate is enzymatically active, we monitored the formation of dsRNA from an ssRNA template using a well-established fluorescence-based assay in which the fluorescent dye SYTO9 binds specifically to dsRNA and fluoresces when excited at 485 nm (*54, 55*) using a Yokogawa CV8000 high throughput spinning disk confocal microscope. Analysis of time-lapse images shows an increase in fluorescence intensity (green) within the condensates over time, suggesting the formation of dsRNA. To ensure that these condensates contain RdRp and are not formed by the dsRNA alone, as dsRNA also has the propensity to form phase separated condensates, we performed the polymerization assay using Atto-647N labeled RdRp (1%) and unlabeled RdRp (99%). The result shows the colocalization of RdRp and dsRNA in the condensates, indicating that the RdRp is active within the condensates (Fig. S7).

### Liquid-to-solid transition triggers the formation of fibrils and inhibits HuNoV GII.4 RdRp activity

To study the dynamics of protein molecules during the maturation of the RdRp condensates, we monitored the material properties of the condensates for 12h using FRAP. In initial experiments, the FRAP data revealed a complete fluorescence recovery (98 %) with a half-life (t_1/2_) of 53.3 ± 2.1 s, which decreased substantially with time (Fig. S8A and B). These condensates retained liquid-like properties for about 4 h and then transformed into gel-like or solid-like condensates. The reduction in the mobility of protein molecules within the condensates suggests that, over time, these condensates change from liquid-like to solid-like, likely driven by the aggregation of proteins. We investigated whether changing the fluidity of the condensates had any effect on the enzymatic activity of RdRp molecules. As described in the above section, we performed the RNA polymerization assay of RdRp post 12 h of incubation at 37 °C. The results indicate that the reduction in the condensate’s fluidity is proportional to the enzymatic activity (Fig. S8C). Negative staining transmission electron microscopy of the phase separated samples after 12 h of incubation at 37 °C showed fibrillar structures of varying lengths (Fig. S8D). Surprisingly, these fibrils have amyloid-like characteristics, such as binding to amyloid-specific dyes thioflavin T (ThT) (Fig. S8E) and Congo red (CR) (Fig. S8F).

### GII.4 infected HIE cells show RdRp containing phase separated condensates

To investigate the formation of these condensates in the context of virus infection, we infected HIEs with GII.4 HuNoV and performed immunofluorescence microscopy using anti-RdRp antibody at different time points post-infection. The microscopy images show that RdRp was initially diffuse and, with time, started forming discrete spherical condensates (Fig. 5A), which gradually increased in size. Since these virus-infected cells needed to be fixed and permeabilized for immunolabeling, we could not perform live cell imaging, including FRAP and fusion experiments. In lieu of this, we examined the sensitivities of these condensates towards the aliphatic alcohol 1,6 HD. The infected cells (24h post-infection) were treated with 5% (v/v) 1,6 HD (added to the cell culture medium) for 5 min and fixed with methanol. The cells were dual immunolabeled with anti-RdRp antibody and anti-VP1 antibody (to detect the level of infection). The confocal images of the 1,6 HD treated HIEs exhibit a reduction in the number of visible condensates compared to the untreated cells, indicating the dissolution of these condensates (Fig. 5B), suggesting their liquid-like nature. To visualize the condensate formation in live cells, we investigated if a recombinant eGFP tagged GII.4 RdRp forms condensates with liquid-like properties upon transfection in HEK293T cells. Confocal microscopy imaging of the cells 24h post-transfection showed round condensates formed by the RdRp (Fig. S9A). In contrast, cells expressing eGFP alone as a negative control exhibit a diffused distribution of eGFP (Fig. S9A). The condensates formed by eGFP-RdRp showed recovery after photobleaching (Fig. S9A and B) and condensate fusion (Fig S9C). Further, these condensates undergo dissolution upon adding 1,6 HD, and they reappear after its removal, indicating their reversible and liquid-like nature. Considering the potential cytotoxic effects of 1,6 HD (*17*) and its limitations (*57*) in the study of LLPS, we also performed the same experiments using 1,2-propane diol or propylene glycol (PG), which showed the same results (Fig S9D). Together, these findings provide evidence that the observed condensates in the virus-infected HIEs harboring RdRp are formed via LLPS.

**Figure 5.**
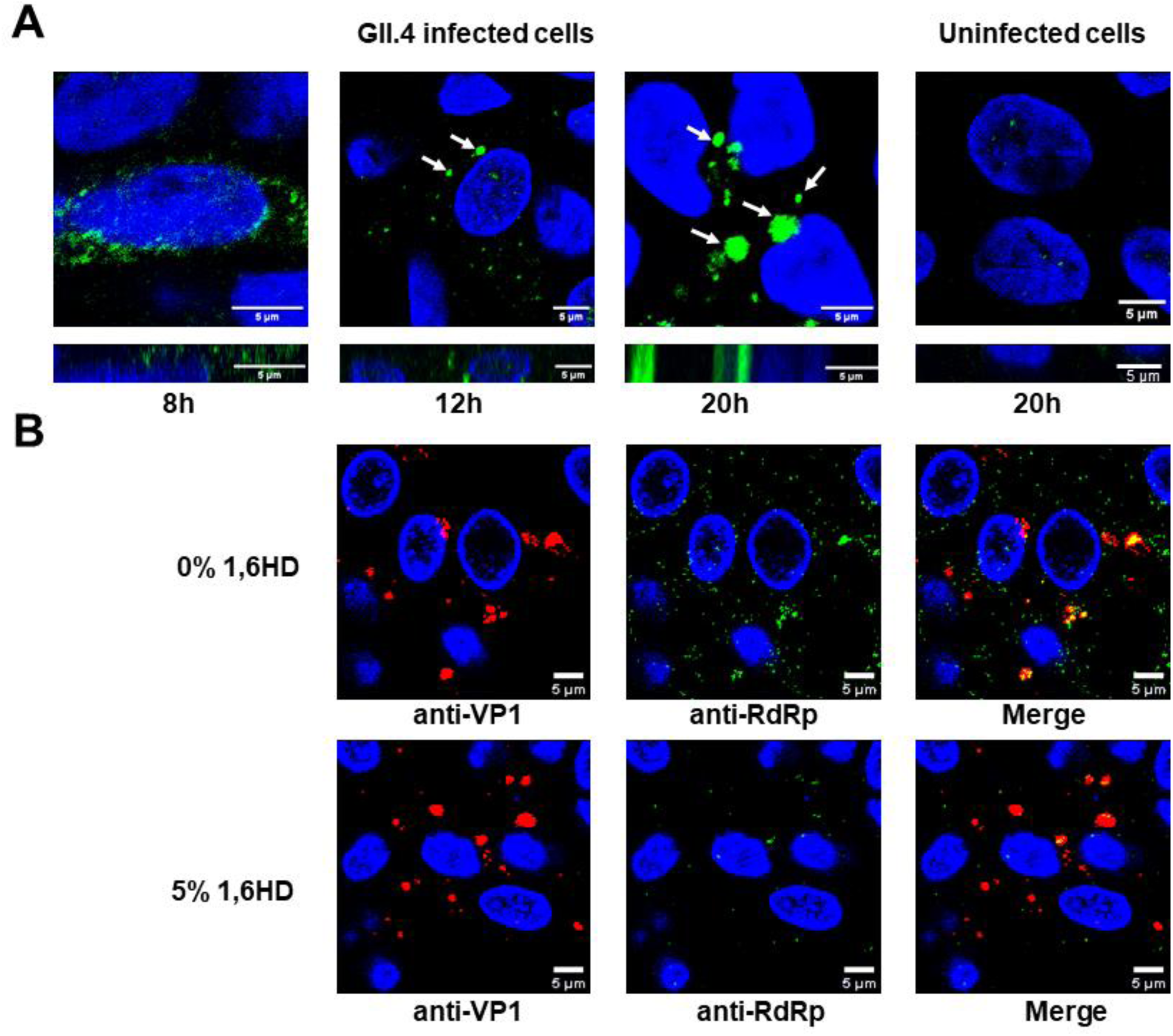
RdRp forms liquid-like condensates in GII.4 infected HIEs. **(A)** Confocal microscopy images of GII.4 HuNoV-infected HIEs (TCH 12-580) at different time points post infection showing condensate formation detected by anti-RdRp antibody (green). Uninfected HIEs were used as negative control. Red arrows indicate the position of representative condensates. **(B)** Effect of 1,6 HD on the condensates formed in the GII.4 HuNoV-infected HIEs (BCM 16-2) suggesting their liquid like nature.

### Condensates in virus-infected HIEs are RFs

To determine if the condensates observed in the virus-infected HIEs are the sites for viral replication, we examined the colocalization of the RdRp with the replicative intermediate dsRNA. Virus-infected HIEs were fixed, immunostained by antibodies against both GII.4 RdRp and dsRNA and imaged using confocal microscopy. Colocalization of dsRNA with RdRp in the condensates (Fig. 6A) was found, suggesting that these condensates are RFs. We further examined the condensates for the presence of VPg, another critical component of the replication machinery (*44*), using antibodies against both RdRp and VPg (Fig. 6B). Similar results were obtained when purified RdRp was mixed with ssRNA and VPg (Fig. S10A, B respectively) The images show the coalescence of RdRp and VPg in these condensates. Together, these results strongly suggest that these condensates are sites for viral replication.

**Figure 6.**
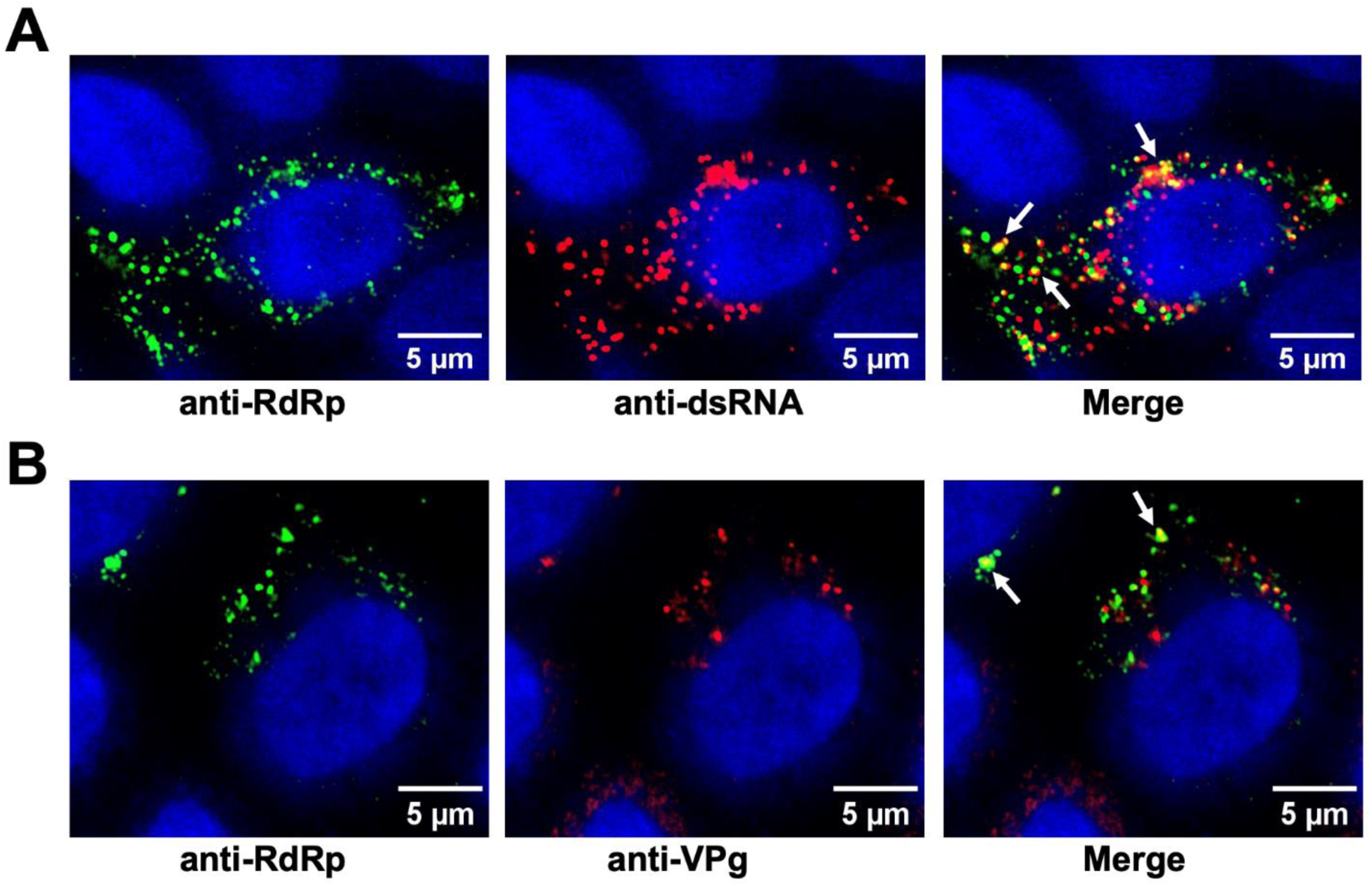
GII.4 RdRp condensates are the sites for viral replication. Confocal images of the GII.4 HuNoV-infected HIEs (BCM 16-2) showing **(A)** Colocalization (yellow) of dsRNA (red) and RdRp(green) detected by respective antibodies **(B)** Colocalization (yellow) of VPg (red) and RdRp(green) detected by respective antibodies. In both the panels white arrows indicate the site of colocalization.

### RdRp condensates are associated with markers for Golgi and ER

Previous studies on picornaviruses, murine noroviruses, and other positive sense RNA viruses have suggested a possible association of the RFs with membranes (*18, 58*),(*59, 60*). To identify the association of any membrane components in the condensates formed during virus infection, we examined the localization of various cellular membrane markers, including *trans*-Golgi body (GalT), *cis*-Golgi body (GM130), ER (calnexin), *cis/medial*-Golgi body (Giantin), and lysosomes (LAMP1) with RdRp marker. In contrast to GalT, which showed clear colocalization with RdRp (Fig. 7A), no colocalization was detected with GM130, calnexin (Fig.s 7B and C), Giantin, and LAMP1 (Fig. S11A and B). These results suggest that HuNoVs recruit membranes derived from late compartments within the secretory pathway, especially *trans-*Golgi bodies, with some specificity. We also investigated if any of the HuNoV nonstructural proteins with a membrane binding domain, such as p41 (NS3), which is an NTPase and viral helicase and considered to be critical for viral replication (*61*), is associated with the RdRp condensates. We performed these experiments in HEK293T cells by co-transfecting RdRp-eGFP and p41-iRFP. The confocal imaging of these cells shows that p41 co-localizes with RdRp, suggesting that RdRp condensates can sequester membrane proteins involved in viral replication (Fig. S11C).

**Figure 7.**
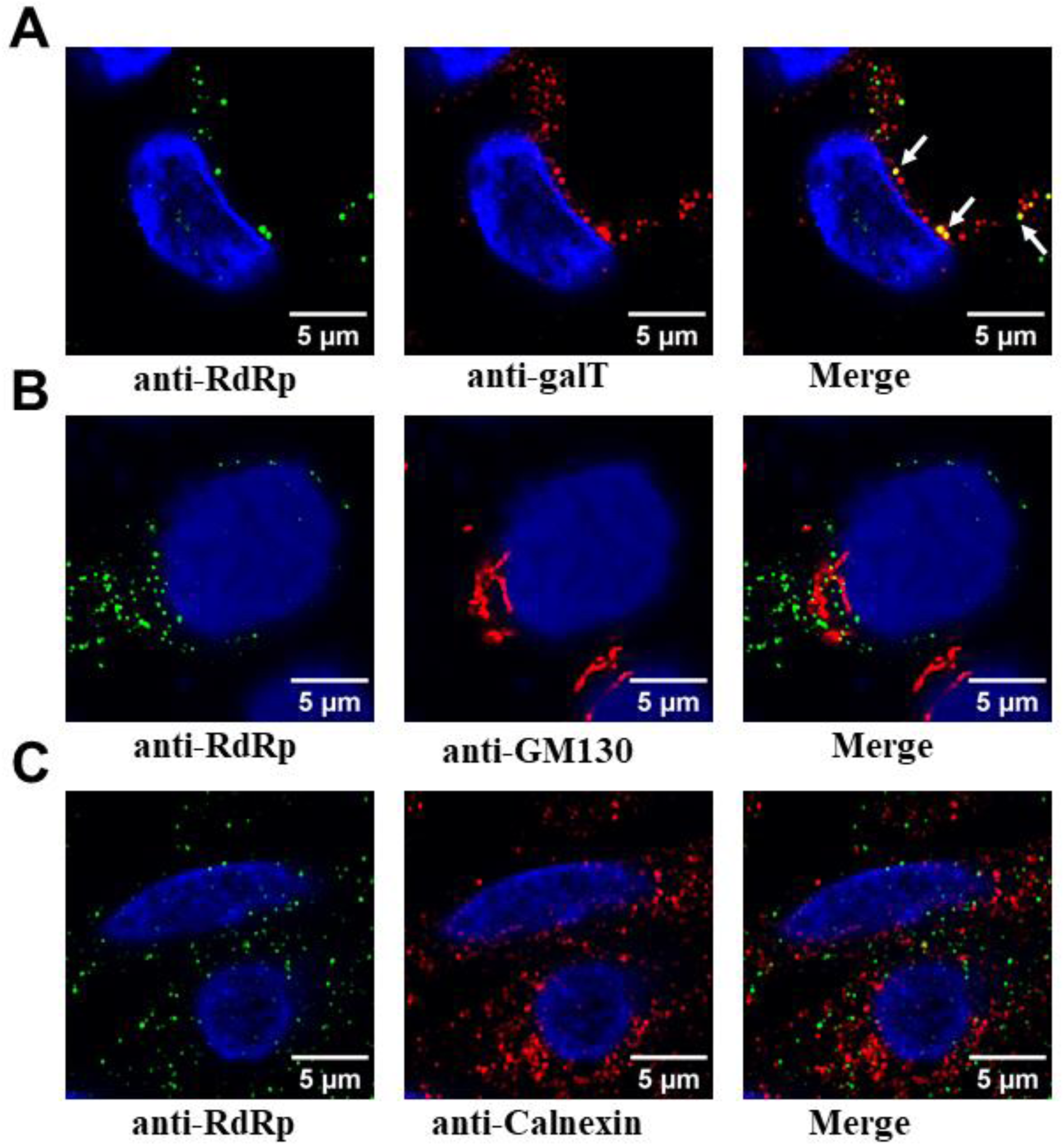
Association of RdRp condensates with membrane markers. GII.4 HuNoV-infected HIEs (BCM 16-2) were fixed and dual labeled with markers for RdRp (green) and various cellular membrane markers (red) **(A)** for the *trans*-Golgi (GalT) **(B)** *cis* Golgi (GM130) **(C)** ER (calnexin).

### Amyloid formation in virus-infected HIEs

To investigate if amyloid-like fibrils, as observed in our in vitro experiments (Fig. S8), are also formed in the context of viral replication, we stained the GII.4 HuNoV infected HIEs with the Thioflavin S dye commonly used to stain amyloids formed inside cells and performed microscopy imaging 48 hours post-infection. Consistent with our in vitro observations, the images show amyloid formation within RdRp condensates (Fig. S12). Such a transformation from liquid-like to solid-like amyloid fibrils might provide a mechanism to regulate viral replication.

## Discussion

In addition to their demonstrated role in regulating various cellular functions, biomolecular condensates formed via LLPS are strongly implicated in forming viral RFs/inclusion bodies (*13, 15–17*). For example, in the case of negative-sense RNA viruses, the involvement of LLPS in the formation of inclusion bodies, which are the sites for viral genome replication, has been extensively studied (*13, 15, 16, 62*). The viroplasms formed during rotavirus infection, the genome replication and sub-particle assembly sites, are considered biomolecular condensates formed by the nonstructural proteins NSP5 and NSP2 (*17*). In the case of SARS-CoV-2, the N protein undergoes LLPS and sequesters components of the viral replication machinery (*63*). Although RFs during HuNoV infection are not yet described, distinct RFs are observed in the case of murine NoV infection (*58*). However, the involvement of LLPS in these RFs has not been explored. In this study, we show that purified RdRp of GII.4 HuNoV undergoes phase separation to create liquid-like, dynamic condensates that contain a protein-rich, dense, yet highly mobile environment. Our studies further suggest that these condensates observed in HEK293T cells and in GII.4 HuNoV-infected HIEs are spherical, grow over time, and are sensitive to 1,6 HD or PG treatment, suggesting their liquid-like nature. We posit that these condensates are the sites for viral replication, as they contain the replicative intermediate dsRNA and necessary replication components RdRp and VPg. Bioinformatics analysis shows that RdRps of almost all the norovirus genogroups have a high propensity to form LLPS (Fig. S12), suggesting that LLPS formation by RdRp may be a common phenomenon in the Norovirus genera.

### Unique aspects of HuNoV GII.4 RdRp induced LLPS – initiating a hub for viral replication

Typically, cellular or viral proteins require additional client proteins, crowding agents, or RNA to phase separate under physiological conditions. For example, up to 50 μM of the SARS-CoV-2 N protein does not phase separate, but adding 1 μM poly U promotes the phase separation (*23*). Similarly, 35 μM NSP5 of rotavirus does not phase separate by itself; but requires the presence of NSP2 or poly-arginine for phase separation (*17*). In contrast, GII.4 RdRp does not require any other viral protein or RNA for phase separation. It can phase separate at 5 μM by itself and at 500 nM by adding a molecular crowding agent. Another interesting result, which may have implications for viral replication, is that RdRp undergoes LLPS at pHs ranging from 5 to 9., Also at pH 5, the minimum concentration of RdRp to form LLPS is reduced from 5 μM to 500 nM. This property may be of relevance considering that virus infection can alter the intracellular pH (*50–52*). The ability of RdRp to readily induce LLPS under various protein concentrations and buffer conditions while remaining enzymatically active supports the idea that during viral infection it likely initiates the formation of a sustainable platform for efficient replication.

### HuNoV RFs are liquid-like condensates

Our experiments in HuNoV-infected HIEs strongly suggest that the condensates observed during viral infection with liquid-like properties are the sites for viral replication as both the replicative intermediate dsRNA and the viral nonstructural protein VPg colocalize with RdRp in these condensates. VPg is a critical protein involved in viral replication and translation of viral proteins. A recent study on murine NoV suggested that apart from its priming function, VPg can physically interact with RdRp and enhance its polymerization activity (*44*).

A question that has to be considered while discussing the RFs as phase separated condensates in positive-sense RNA viruses is whether they are associated with the membrane or membrane components. Although not unequivocally demonstrated (*64, 65*), earlier studies on picorna-like viruses (*66*) and murine noroviruses have suggested that RFs are associated with intracellular membranes and vesicular structures (*18, 58–60*). The formation of LLPS as a potential hub or a driving force for forming the RFs does not exclude the incorporation or the association of membrane-anchored proteins and lipids. Recent studies have implied such an association could reduce the critical concentration for inducing LLPS and further for forming microdomains within the membrane-associated condensates (*67–69*). Our studies with membrane markers in HuNoV infected HIEs show colocalization of RdRp with the *trans-Golgi* marker galT in the condensates. The markers for other membrane components like *cis*/*medial*-Golgi (GM130), ER (calnexin), cis/medial-Golgi body (Giantin), and lysosomes (LAMP1) show very little or no association with RdRp in the condensates. In positive-sense RNA viruses, particularly the picornavirus superfamily, the protein (s) involved in the biogenesis of the RFs usually contain a domain that associates with the membrane or interacts with another viral protein that anchors it to a cellular membrane, thereby creating a vesicular (membrane-associated) replication organelle (*18, 19*). Our results in HEK293T cells show the sequestration of p41 (NS3), a membrane-anchored protein with RNA helicase activity (*61*) to the RdRp condensates, leading to the possibility that p41 acts as a bridge between RdRp and the membranous component of replication complex. Further studies are required to address the possible co-existence of phase separated condensates in the context of vesicular structures in the replication of HuNoV and other picorna-like viruses.

### A mechanism to resolve the translation-replication conundrum?

The formation of the RFs via LLPS in positive-sense RNA viruses may have a distinct advantage, as has been recently suggested (*70*). In positive-sense RNA viruses, genomic RNA serves as the basis for both translation by the ribosomes, which is processed from the 5’ to the 3’ end, and replication by the RdRp from the 3’ to the 5’ end. These two processes running in opposite directions pose a conundrum for the replication of these viruses, as previously recognized (*71*). The process of phase separation could provide an elegant solution for resolving this conflict by isolating the newly synthesized RdRp capable of sequestering the replication components from interfering with ribosomal translation. In the case of HuNoVs, RdRp, as we suggest, triggers the biogenesis of phase separated RFs; however, in other positive-sense RNA viruses, it is quite possible that another viral protein that is essential for replication could trigger this process by undergoing LLPS, and then recruit the RdRp into the condensates.

### Liquid-to-solid transition as a regulatory mechanism

Another common phenomenon of biomolecular condensates is the transition from liquid to solid state over time. The FRAP analysis of the GII.4 RdRp condensates over time show a lower diffusion rate of the RdRp molecules in the condensates. Consistent with this observation, TEM imaging of the RdRp incubated at 37 °C overnight demonstrated the formation of fibrils. Interestingly, these fibrils bind to amyloid-specific dyes such as ThT and CR, indicating their amyloid-like nature. A similar transition is also observed in virus-infected HIEs. Interestingly, RdRps of several picornaviruses also form similar fibrillar structures (*72, 73*), although whether they exhibit amyloid-like properties is still unknown. We also observed that under these conditions, the enzymatic activity of RdRp is considerablyreduced, supporting the notion that the transition from liquid to solid state is plausibly a mechanism for regulating replication. This could be the reason why all of the puncta are not dissolved by 5% 1,6HD, and do not show colocalization of RdRp with dsRNA and VPg (Figs 5B and 6), as some of them may have transitioned to the solid-like phase. Another possibility is that these amyloid-like fibrils may have a role in inducing cell toxicity, as shown in the case of amyloid-like fibrils formed by the NS11 protein of SARS-CoV-2 (*74*).

In summary, our study reveals a previously unknown property of HuNoV RdRp in forming biomolecular condensates as platforms for genome replication. In addition to exhibiting all the properties for inducing the formation of LLPS in vitro, we have shown that RdRp also readily undergoes LLPS in a cellular environment that resembles the condensates observed in virus-infected HIEs strongly implicating RdRp in initiating and sustaining the viral RFs during HuNoV infection. Our bioinformatics analysis suggests that this indeed may be a common phenomenon in most noroviruses. The implication of RdRp in the biogenesis of RFs also opens new targets for designing antivirals for HuNoV infections, which remains a serious threat in children and immunocompromised patients. Formation phase separated condensates as replication hubs may perhaps be a necessary mechanism in the positive-sense RNA viruses, whether initiated by the RdRp or other viral proteins, to effectively segregate the otherwise conflicting translation and replication events.

## Materials and Methods

### Cloning, expression, and purification of proteins

Codon-optimized RdRp and VPg genes of HuNoV GII.4 Sydney strain (GenBank: AFV08794.1) were synthesized and cloned in the pET28a vector for bacterial expression by Genscript U.S.A. Inc with an N-terminal 6x His-tag to aid in the purification of both proteins. For mammalian expression HuNoV GII.4 Sydney strain (GenBank: AFV08794.1) RdRp was cloned with eGFP on C-terminal, VPg with mCherry on N-terminal, p41 with iRFP on N-terminal in pcDNA3.1(+) by Genscript U.S.A. Inc. The N-terminal truncation, i.e., GII.4 RdRp-NΔ51 for bacterial expression were also generated by Genscript U.S.A. Inc. For the over-expression of proteins, the plasmids containing the desired genes were transformed into *E coli* BL21 (DE3) cells. Protein expression was induced by the addition of 0.3 mM isopropyl β-D-1-thiogalactopyranoside at OD_600_ ∼0.6. Cells were allowed to express the protein at 18 °C for 16h with constant shaking at 200 rpm. Cells were harvested by centrifugation at 4000xg for 30 min at 4 °C and the pellet was re-suspended in lysis buffer (20 mM Tris-HCl pH 7.5, 250 mM NaCl, 5 mM imidazole, benzonase nuclease, and a cocktail of protease inhibitors) followed by cell disruption using a microfluidizer. The supernatant was collected after centrifugation at 18,000xg for 45 min at 4 °C. The protein was purified by affinity chromatography using Ni-NTA agarose (Qiagen) resin followed by his tag cleavage by TEV protease and gel filtration using Superdex 200 Increase 10/300 GL column (G.E. Healthcare) in 20 mM Tris-HCl pH 7.5, 150 mM NaCl and 2 mM DTT. All the target proteins GII.4 RdRp, GII.4 RdRp-NΔ51, and GII.4 VPg were purified using the above-mentioned protocol to a purity of >95% as judged by SDS-PAGE and Coomassie blue stain.

### Analytical ultracentrifugation

AUC experiments were carried out as previously described (*75*) with minor modifications. Briefly sedimentation velocity experiments of 10 μM GII.4 RdRp at pH 5.5, 7.5, 9.0 were carried out using a Beckman-Coulter XL-A analytical ultracentrifuge with a TiAn60 eight-hole rotor and two-channel epon centerpieces (12 mm) at 40,000 rpm at 22 °C. Absorbance scans were recorded at 280 nm at 1 min intervals. Experiments were carried out three times.

### Protein crystallization

Crystallization screenings for GII.4 RdRp (10 mg/ml) were carried out by hanging-drop vapor diffusion using the Mosquito crystallization robot (TTP LabTech) and imaged using Rock Imager (Formulatrix) at 20 °C. The protein was crystallized under the conditions 0.3 M Sodium Iodide, 0.1 M Sodium Acetate: HCl, pH 4.5, 22.5 % (v/v) PurePEGs Cocktail. Crystals were flash-frozen directly in liquid nitrogen.

### Structure determination and refinement

X-ray Diffraction data for GII.4 RdRp was collected on APS beamline17-ID. Diffraction data were processed with IMOSFLM as implemented in the CCP4 suite (*76*). The structure of NV RdRP (PDB ID: 4LQ3) was used as a search model for molecular replacement (MR) using PHASER. The atomic model, including the side chain atoms, was then subjected to iterative cycles of refinement using PHENIX and further model building using COOT based on the difference maps (*77, 78*). Data collection and refinement statistics following the final refinement cycle are given in Supplementary Table 1.

### SEC-SAXS data collection and processing

The SEC-SAXS data for HuNoV GII.4 RdRp (5mg/ml, monomer) was collected at the SIBYLS beamline 12.3.1 at the Advanced Light Source facility. The X-ray wavelength was fixed to λ = 1.127 Å, and the distance between the sample and the detector was set to 2100 mm, resulting in scattering vectors (q) spanning from 0.01 to 0.4 Å−1. RdRp (60 μl) sample was prepared in running buffer composed of 20 mM HEPES at pH 7.5, 150 mM NaCl, and 0.1% β-mercaptoethanol. The Shodex KW803 column was equilibrated with the same running buffer, running at a flow rate of 0.5 ml/min. Each sample, consisting of 55 μl, was injected into the SEC column, and X-ray exposures lasting 3 seconds were continuously recorded for over 30 minutes. Buffer subtraction was performed using SCÅTTER (https://bl1231.als.lbl.gov/scatter/), and the resulting curves were merged for subsequent Guinier analysis, Dimensionless Kratky plots, and P(r) plot computations.

### Solution structure modeling

The crystal structure of RdRp (PDB ID: 8TUF) was used as a template for rigid body modeling. Minimal molecular dynamic simulations were performed on the flexible N-terminal regions within the structure to investigate the conformational space adopted by RdRp using BILBOMD (*47*). This approach involved storing a single conformational state approximately every 100 ps, yielding a total of 10,000 conformers varying in Rg and Dmax values for subsequent SAXS fitting and multistate validation (*79*). Initially, scattering profiles from all 10,000 models were computed, followed by genetic algorithm-selection operations to refine the selection to the most suitable models. Finally, experimental scattering profiles from each construct were compared with theoretical scattering profiles of the chosen atomistic models generated by BILBOMD, utilizing FOXS (*80*).

### Light scattering measurements

Light scattering measurements of GII.4 RdRp was carried out on a FlexStation3 plate reader (Molecular Devices) using a 96-well flat bottom plate (Fisher Scientific) by measuring absorbance at 350 nm at 37 °C at a regular interval of 2 min. Different concentrations of GII.4 RdRp (0.5, 1, 5, 10, 25 μM) were screened at varying concentrations of PEG3350 (0, 5, 10, 20%), pH (5.0, 6.0, 7.5, and 9.0) and salt (NaCl) (0, 150, 300, 500, and 750 mM). Experiments were carried out three times.

### Fluorescence labeling of proteins

For microscopic visualization of condensates, GII.4 RdRp and GII.4 RdRp-NΔ51were labeled with Atto-488 and GII.4 VPg was labeled with Atto-647N dye. For labeling an Atto 488 protein labeling kit (Sigma-Aldrich, 38371) and an Atto 647N protein labeling kit (Sigma-Aldrich, 76508) were used. The manufacturer’s provided protocol was used. Briefly, 8 mg/ml RdRp, RdRp-NΔ51, and 6 mg/ml VPg were dialyzed in sodium bicarbonate buffer solution, pH 9.5 (provided with kit).

Then the protein solution was transferred to a vial containing the reactive dye dissolved in 20 μl sodium bicarbonate buffer and incubated the reaction mixture for 2 h at room temperature while gently stirring and protecting the vial from light. For the separation of protein-dye-conjugates from free dye, the manufacturer’s provided gel filtration columns were used to achieve good separation of the protein-dye-conjugates from the excess free dye (size exclusion limit: 5 kDa). The protein-dye-conjugates were eluted in phosphate buffer, pH 7.5 provided with the kit.

### Confocal microscopy

Protein mixtures of 1% labeled and 99% unlabeled proteins were made at a concentration of 10 μM. An aliquot (15 μl) of the protein mixture was spotted onto 15 mm depression slides and sandwiched with an 18 mm coverslip followed by sealing of the edges of the coverslip with commercially available nail polish and allowed to phase separate at 37 °C for 45 min. The condensates were visualized by Nikon A1R-s confocal microscope with 40X-air objective and 1 Airy unit aperture.

Different concentrations of GII.4 RdRp (0.5, 1, 5, 10, 25 μM) were screened at varying concentration of PEG3350 (0, 5, 10, 20 %), pH (5.0, 6.0, 7.5, and 9.0) and salt (NaCl) concentration (0, 150, 300, 500, and 750 mM) using a Yokogawa CV8000 high throughput spinning confocal imaging platform with 60x/1.0 water immersion objective lens. Four fields of views with multiple Z planes were captured. Experiments were carried out three times.

### Far-UV circular dichroism (CD) measurements

Far-UV CD measurements of RdRp and RdRp-NΔ51 were carried out using Circular Dichroism Spectroscopy (J-1500, Jasco), using a 1-mm path-length quartz cuvette. Data were recorded using a final protein concentration of 8 μM proteins in 20 mM sodium phosphate buffer pH 7.5. The spectra were averaged over three scans and were blank subtracted. The spectra were then processed using Spectra Manager™ CDPro (Jasco) and plotted using Origin 2018.

### Negative-stain transmission electron microscopy (EM)

GII.4 RdRp (10 μM, 50 µl) was incubated at 37 °C at a constant shaking at 200 rpm for 20 s after every 2 min for a period of 12 h. For preparing negatively stained grids, 3 μl of the GII.4 RdRp (diluted 10 times) was adsorbed onto a glow-discharged grid carbon-coated grid (CF200-CU; EMS) for 60 s. The grid was blotted with Whatman’s filter paper 541 and washed twice with deionized water with intermittent blotting. The grid was further stained with three drops of 2 % uranyl acetate with intermittent blotting to remove the residual stain. The grids were air-dried and imaged at room temperature using a JEM 2100 transmission electron microscope operated at 200 keV (JEOL) equipped with a LaB6 filament, 3kx4k Direct Electron Detector (DE12) and a Gatan 4kx4k CCD at the cryo-EM core facility, Baylor college of Medicine. The images were recorded at 15k using a low-dose procedure and a defocus range of −1 to −4 μm.

### GII.4 HuNoV infection of HIE monolayers

Five day differentiated monolayers were infected with 1 x 10^7^ GE of GII.4 stool filtrate (TCH 12-580 or BCM 16-2) in CMGF(-) for 1 h at 37 °C. After washing two times with CMGF(-), the HIEs were incubated at 37 °C for different time points (8-20 h). The cells were fixed with 100% chilled methanol for 20 min at room temperature. The cells were incubated with 1:150 dilution of rabbit anti-RdRp (*81*) in 5 % BSA PBS at 4 °C overnight. The cells were washed three times with PBS containing 0.1 % Triton X-100 and incubated with goat anti-rabbit 488 (#A-11008, ThermoFisher Scientific) overnight at 4 ° C to visualize the viral proteins. The cells were washed three times, and the nuclei were stained with 4, 6-diamidino-2-phenylindole (DAPI) (300 nM) followed by subsequent Z-stack images captured using a Nikon A1 confocal microscope. For treatment with 1,6 HD, the cells post-infection 20 h were treated with 5% 1,6 HD which was added to the cell culture medium for 5 min and fixed with 100% chilled methanol as described above. The cells were incubated with 1:150 dilution of rabbit anti-RdRp (*81*) and 1:500 dilution of guinea pig VP1antibody in 5 % BSA PBS at 4 °C overnight and washed and imaged as described above. For dual leveling experiment of RdRp with dsRNA, VPg and membrane components, the antibodies used are mouse anti-dsRNA (Millipore Sigma), anti-VPg (in house produced), anti-GalT (Abcam), anti-GM130 (Abcam), anti-calnexin (Abcam), anti-Giantin, and anti-LAMP1(Santa Cruz Biotechnology). For thioflavin S (THS) staining the cells were incubated with 0.05% THS (dissolved in 50% ethanol/water) for 15 min at room temperature after the secondary. The cells were washed twice with 50% ethanol in water for 20 min each followed by one wash with 80% ethanol in water for 20 min.

### Mammalian cell line, expression and treatment with aliphatic alcohols

HEK293T (ATCC, CRL-3216) (2 x 10^5^ cells per well) were seeded in µ-Slide 8-well glass-bottom plates (Ibidi) in DMEM media and incubated at 37 °C for 24 h. Cells were then transfected with respective plasmids and incubated at 37 °C. After 24 h, the cells were stained with Hoechst 33342 diluted in FluoroBRITE DMEM at a concentration of 1 mg/mL and incubated for 30 min at 37 °C. For treatment with 1,6 HD and PG, live cell imaging of eGFP-RdRp transfected HEK293T cells was performed after 24 h using confocal microscopy. After imaging a suitable area for condensates, the media was replaced with FluoroBRITE DMEM media supplemented with 5% 1,6 HD or 5% PG. The imaging was continued in the same area, observing for dissolution of condensates for 5 min. The media was again replaced with FluoroBRITE DMEM without the respective aliphatic alcohols to observe the reoccurrence of the condensates in the same area. Experiments were carried out three times.

### Fluorescence recovery after photobleaching (FRAP)

FRAP studies were performed using a laser scanning confocal microscope (Nicon-A1R-s) with a 40x air objective. A circular area encompassing an entire condensate was bleached with a 404 nm laser (100 %) for 20 iterations and recovery was recorded for 300 s at 10 s intervals. For in vitro FRAP study, 15 μl RdRp (10 μM) was spotted onto 15 mm depression slides and sandwiched with 18 mm coverslip followed by sealing of the edges of the coverslip with commercially available nail polish and allowed to phase separate at 37 °C. After droplet formation was confirmed, the slides containing the condensates were subjected to FRAP experiments at various time points as indicated in the respective sections. For live cell FRAP, the bleaching of ROI was performed at 70 % laser power for 20 iterations and recovery was recorded for 500 s at 15 s interval. The images were corrected for laser bleaching by selecting a fluorescent region outside the ROI.

### Estimation of light phase and dense phase concentration

The concentrations of the dense phase and light phase were estimated using centrifugation by following a previously described protocol (*56*). For dense phase concentration estimation, 100 μl RdRp (10 μM) was incubated at 37 °C for 45 min and centrifuged at ∼ 25,000 x g at 37 °C for 15 min. After carefully removing the supernatant in several steps, the pure dense phase (∼ 2 μL) was re-suspended in 100 μl of the denaturation buffer (6M guanidine hydrochloride, 20 mM sodium phosphate). The absorbance at 280 nm was recorded using nanodrop. The light phase concentration was estimated from the absorbance of the supernatant at 280 nm.

### Fluorescence-based activity assay for GII.4 RdRp

GII.4 RdRp activity was measured using a real-time fluorescence-based assay, which uses SYTO9, a fluorescence dye that specifically binds dsRNA but not ssRNA template molecules. Reactions were performed in individual wells of black 96-well flat-bottom plates (costar). The standard reaction contained GII.4 RdRp (monomer or higher order oligomer, 1 μM), 20 mM Tris-HCl pH 7.5, 5 mM MgCl2, 2.5 mM MnCl2, 40 μg/mL polyC, 5 U RNAseOUT (invitrogen) and 0.25 μM SYTO9 (Sigma-Aldrich). The reaction was initiated by the addition of 300 μM GTP, and the fluorescence was recorded every 2 min over 60 min at 30 °C using a plate reader FlexStation3 (Molecular devices). For measuring the activity of RdRp after incubation at 37 °C, the incubated RdRp (10 μM) was centrifuged at 20,000xg for 20 min. The supernatant was removed, and the pallet was mixed thoroughly, and activity was measured using the same protocol as described above. To measure the activity of RdRp [(Atto-647N labeled RdRp (1%) and unlabeled RdRp (99%), 1 μM] in the condensates, the same reaction was set up in 96 Well glass bottom plates (Cellvis), and the fluorescence was recorded every 20 min over 180 min at 37 °C using a Yokogawa CV8000 high throughput spinning confocal imaging platform with 60x/1.0 water immersion objective lens. Four fields of views with multiple Z planes were captured.

### Thioflavin T fluorescence assays

10 μM GII.4 RdRp was mixed with 10-fold molar excess of ThT and fluorescence intensity was measured using excitation wavelength of 440 nm and emission wavelength of 482 nm using a plate reader FlexStation3 (Molecular devices) at constant temperature of 37 °C and readings were acquired every 2 min with continuous orbital shaking between reads over a period of 24 h. The top of the each well was sealed using adhesive tape to minimize the rate of evaporation. Three independent experiments were performed in triplicates and the bars represent the standard deviation.

### Congo red binding assay

CR dye solution was prepared by mixing 7 mg/mL dye in and passed through a 0.22 µm filter prior to use. The spectrum of CR (5 μL in 1 mL 20 mM Tris-HCl pH 7.5, 150 mM NaCl, and 5 mM DTT) was recorded between 400 and 700 nm at room temperature (negative control). 10 μM RdRp was incubated with C.R. for 12 h at 37 °C and the spectrum was recorded between 400 and 700 nm a plate reader FlexStation3 (Molecular devices)

## Acknowledgments

This research used resources of the Advanced Photon Source, a U.S. Department of Energy (DOE) Office of Science user facility operated for the DOE Office of Science by Argonne National Laboratory under Contract No. DE-AC02-06CH11357. We acknowledge SIBYLS beamline 12.3.1 at the Advanced Light Source facility for SEC-SAXS data collection. Imaging for this project was supported by the Center for Advanced Microscopy and Image Informatics (CAMII, CPRIT RP170719) and the Integrated Microscopy Core at Baylor College of Medicine (funding from NIH (DK56338, CA125123, ES030285, S10OD030414). We also acknowledge the CryoEM Core at Baylor College of Medicine for all the TEM experiments. We acknowledge the Protein Characterization Unit, Protein and Monoclonal Antibody Production Core, BCM for AUC experiments. GK was supported by the Cancer Prevention and Research Institute of Texas grant RR160029 (to Xiaodong Cheng, MD Anderson Cancer Center).

## Author contributions

Conceptualization: BVVP, SK

Methodology: SK, RA, BVA, GK, SS

Investigation: SK, RA, BVA,GK

Visualization: SK, BVA, GK, FS

Supervision: BVVP, MKE, SEC, FS, JP

Writing original draft: BVVP, SK

Writing review & editing: BVVP, SK, RA, BVA, GK, SS, SEC, MKE, FS, JP

## Data and materials availability

All data needed to evaluate the conclusions in the paper are present in the paper and/or the Supplementary Materials.

**Supplementary Material** includes 1 table, 13 Figures, and 2 videos with legends.

