## Supplementary data for "RNA-dependent RNA polymerase of predominant human norovirus forms liquid-liquid phase condensates as viral replication factories"

**Revised- adp9333**

**This PDF file includes:**

Figs. S1 to S13
Tables S1
Movies S1 to S2

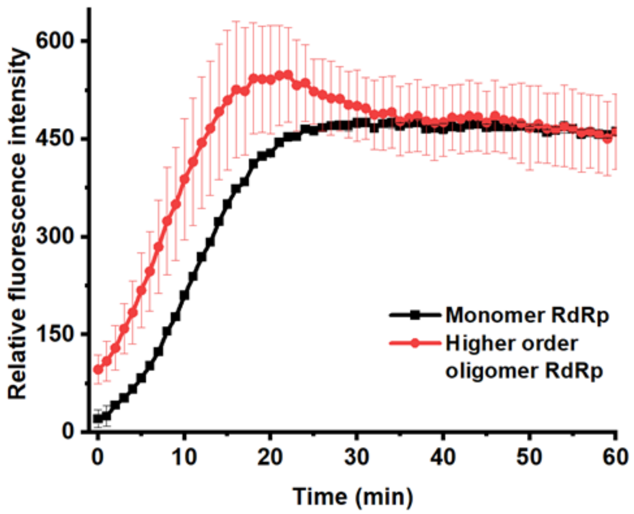


**Figure S1. RNA polymerase activities of monomeric and oligomeric RdRp.** Monomer (black curve) and higher order oligomer fractions of RdRp (red curve) exhibit comparable polymerase activities as measured using SYTO9 dye.

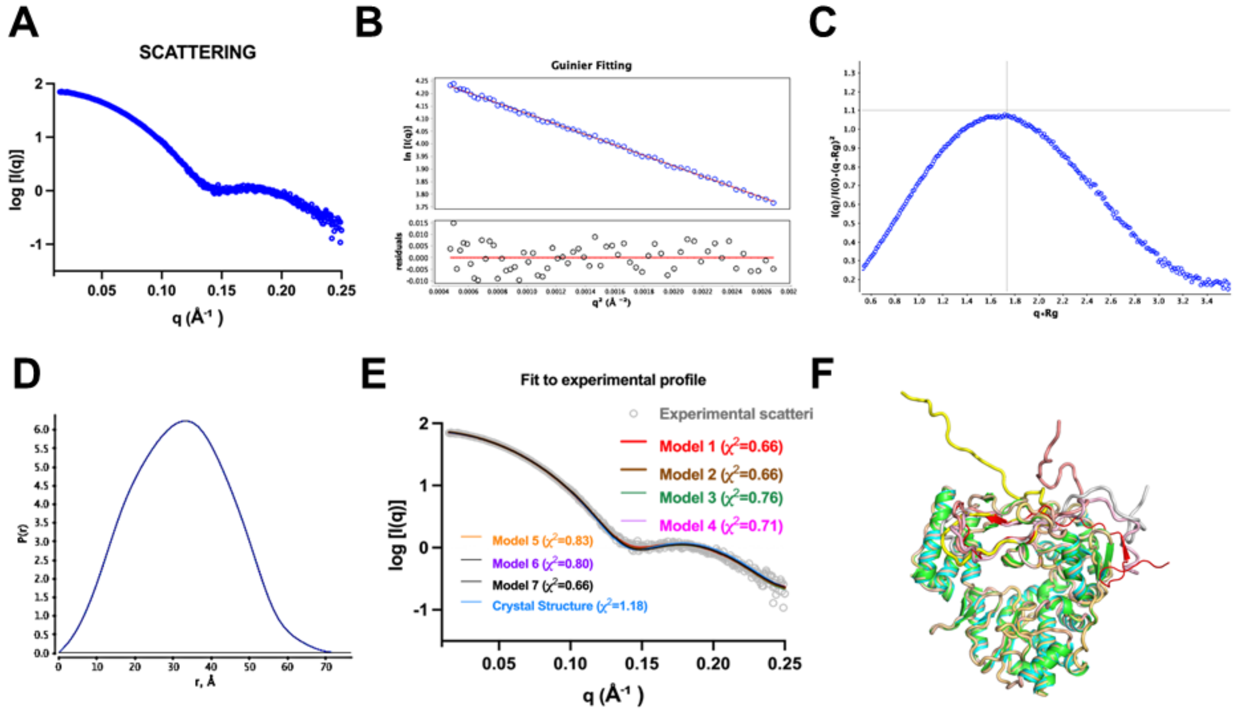

**Figure S2. SEC-SAXS of GII.4 RdRp shows N-terminal region is flexible**. **(A)** Experimental SEC-SAXS scattering intensity profiles. **(B)** Guinier plots from the measured scattering intensity *I*(*q*) as a function of the scattering vector *q* in the low-*q* regions. **(C)** Dimensionless Kratky plots. **(D)** Normalized pair-wise distribution function. P(r) plots computed from the indirect Fourier transformation of measured. **(E)** Comparison of experimental scattering intensity profile with theoretical scattering intensity profile computed from of top eight atomistic BILBOMD model. **(F)** Comparison of the top eight models generated by BILBOMD using the crystal structure of RdRp.

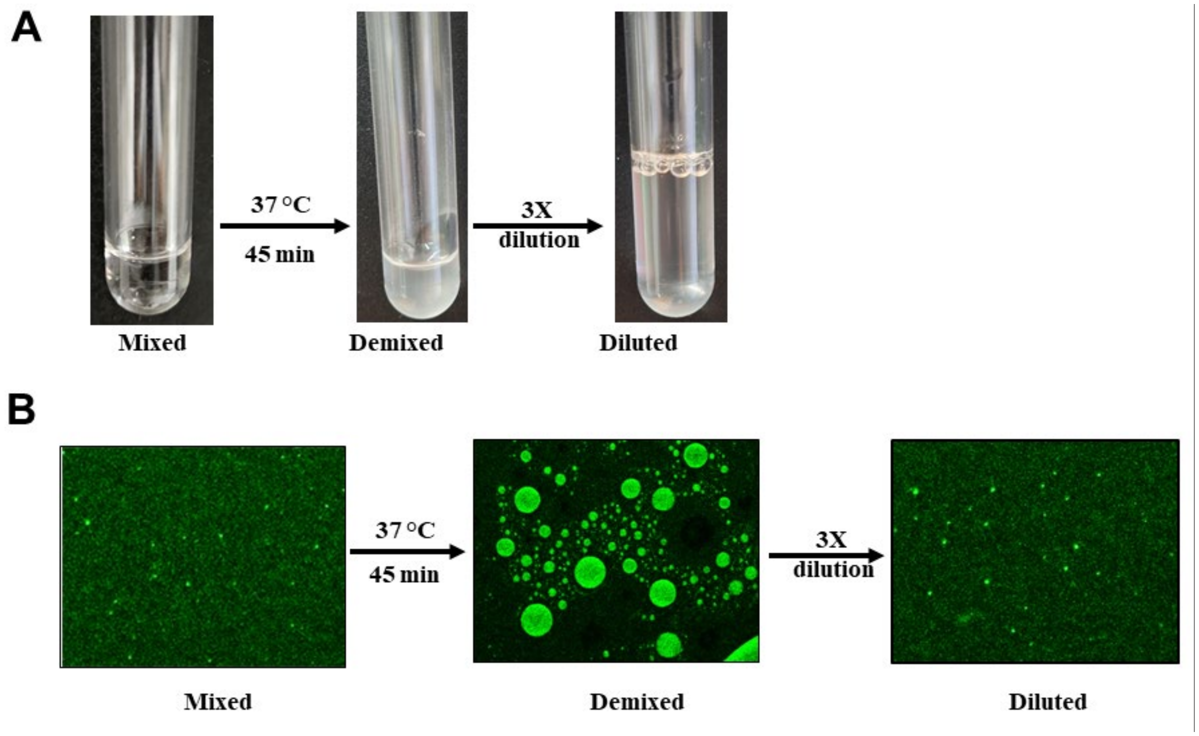


**Figure S3: Reversible nature of RdRp condensates**. **(A)** Visual observation of phase separation of RdRp from a homogeneously mixed phase into a demixed phase upon incubation at 37 °C for 45 min. **(B)** Reversibility of RdRp (green) condensates upon 3X dilution visualized by confocal microscopy.

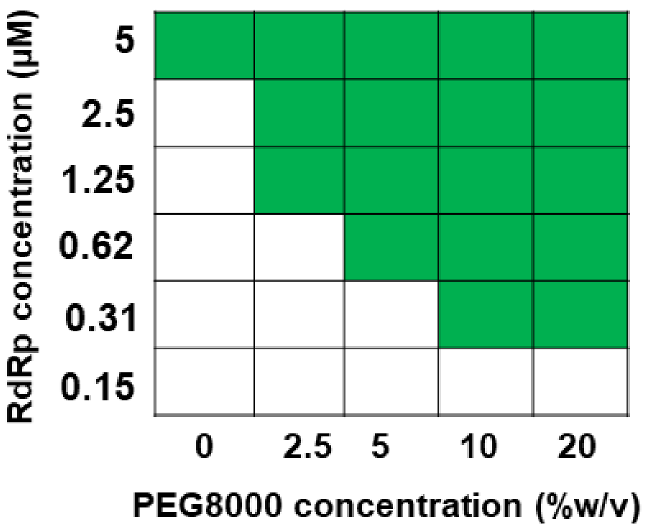


**Figure S4. RdRp phase separates with higher molecular weight crowding agents.** Phase diagram of RdRp condensates at different concentrations of RdRp and PEG8000 which was used as higher molecular crowding agent. The green boxes indicate LLPS (condensate state), and the white boxes indicate no LLPS (soluble state).

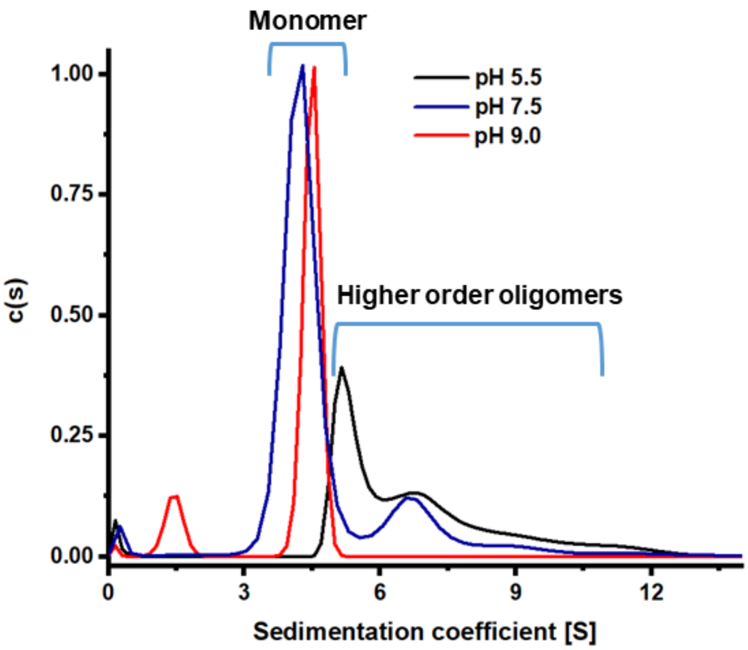


**Figure S5. pH-dependent oligomerization of RdRp**. Analytical ultracentrifugation profiles of RdRp at different pHs shown in different colors as noted in the figure.

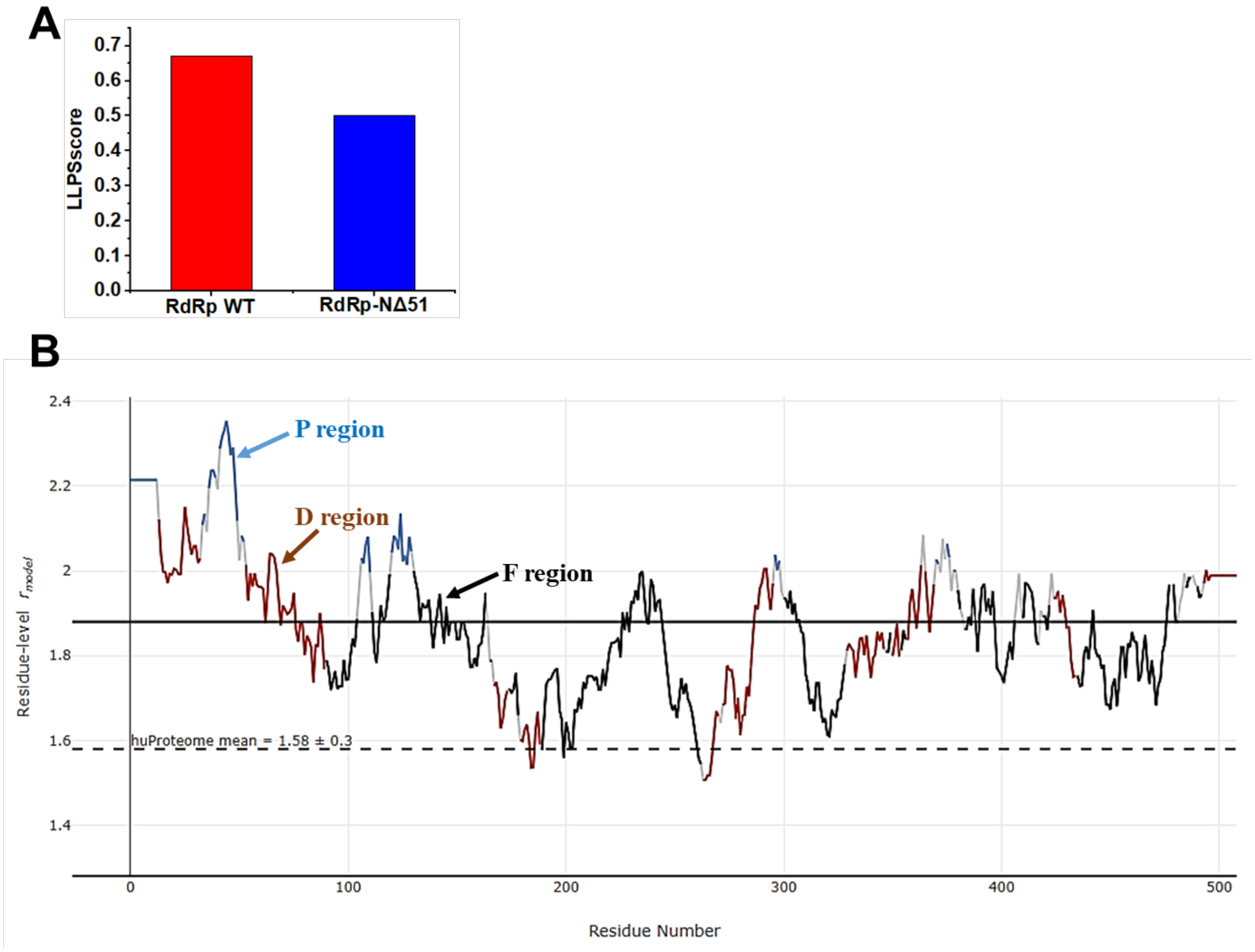


**Figure S6 Bioinformatics analysis shows the N-terminal region is important for LLPS formation. (A)** DeePhase scores for WT RdRp (red) and RdRp-NΔ51(blue) sequences. **(B)** ParSe analysis of WT RdRp. The annotation for different colors in the graph: P regions (blue) are intrinsically disordered and prone to undergo LLPS. D regions (brown) are intrinsically disordered but less likely to undergo LLPS. F regions (black) may or may not be intrinsically disordered but can fold to a stable conformation.

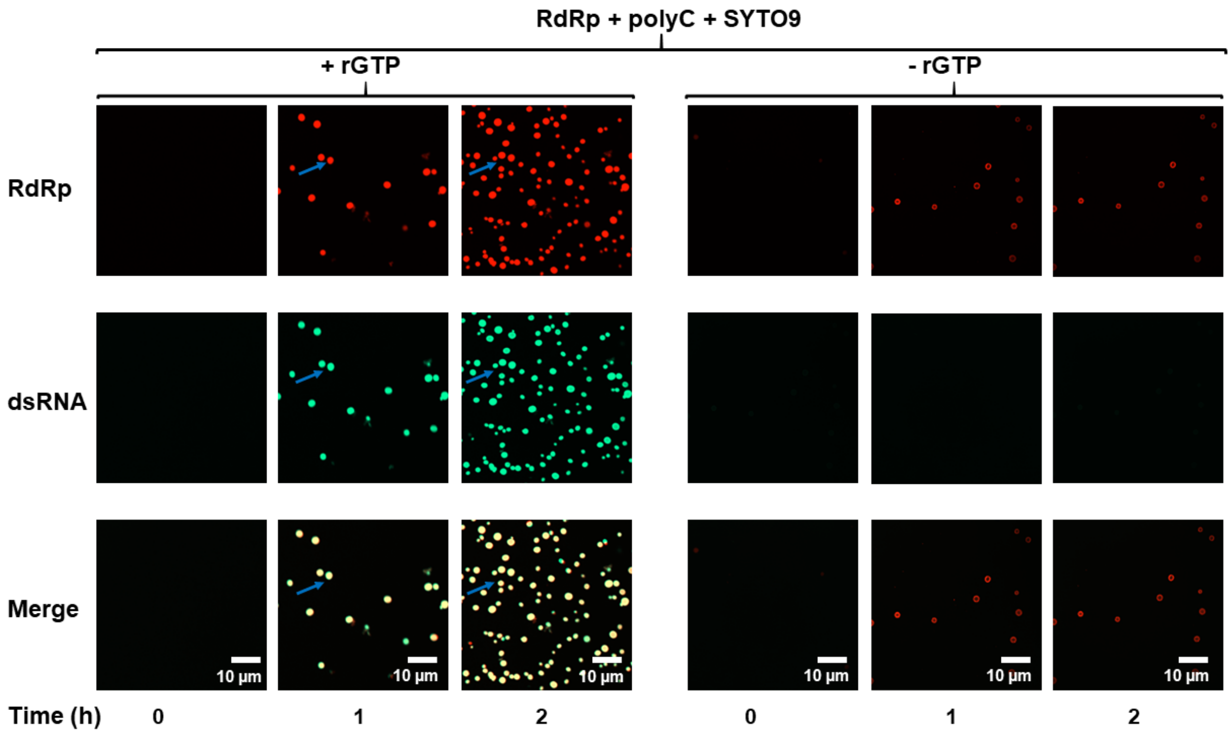


**Figure S7. GII.4 RdRp is enzymatically active in the condensates.** Time-lapse microscopic images of the reaction mixture containing Atto647N-RdRp (red), polyC, rGTP and SYTO9 dye (green) to detect the dsRNA product. The left panel shows the time-dependent increase in the number of RdRp (top) condensates and the synthesis of dsRNA (middle). The merged image (yellow, below) shows the colocalization of RdRP and dsRNA (blue arrows). Corresponding time-lapse microscopic images of the reaction mixture without rGTP in the right panel is shown as a negative control.

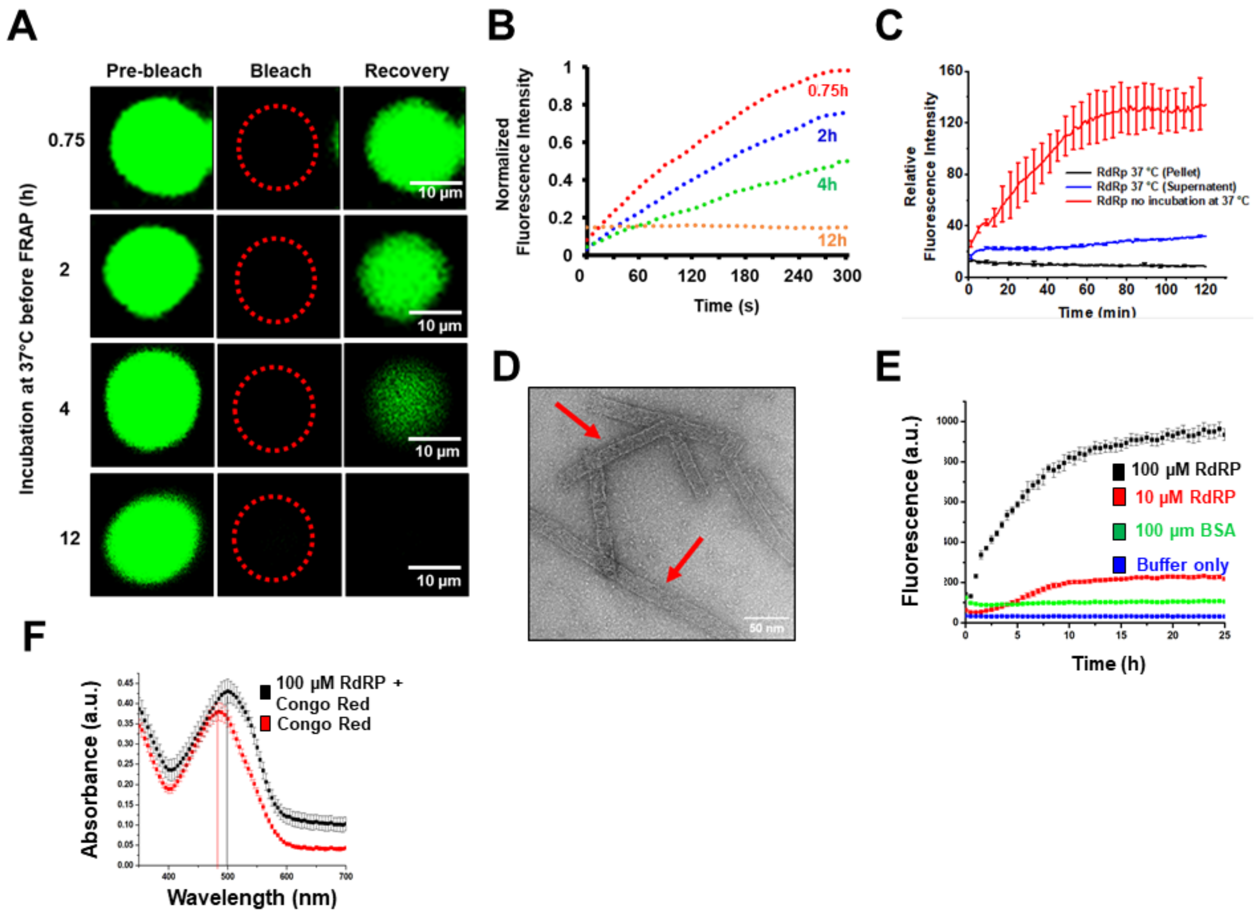


**Figure S8. The dynamics of GII.4 RdRp in the condensates slow down with time leading to the formation of amyloid fibrils. (A)** FRAP of condensates incubated at 37 °C for indicated time points to measure the dynamics of RdRp molecules in the condensates. **(B)** The recovery graph of the FRAP experiments performed in panel A. The results indicate a slowdown in the dynamics of RdRp molecules over time. **(C)** The activity of RdRp after incubation at 37 °C for 12 h was assessed by a RNA polymerization assay using SYTO9 dye. **(D)** Negative stained image of RdRp incubated at 37 °C for 12 h. The red arrows indicate fibrils. **(E)** The kinetics of amyloid fibril formation of RdRp was monitored at 37 °C using a ThT fluorescence assay. **(F)** CR absorbance spectra in the absence and presence of RdRp suggest a red shift in the absorbance maxima of CR in the presence of RdRp.

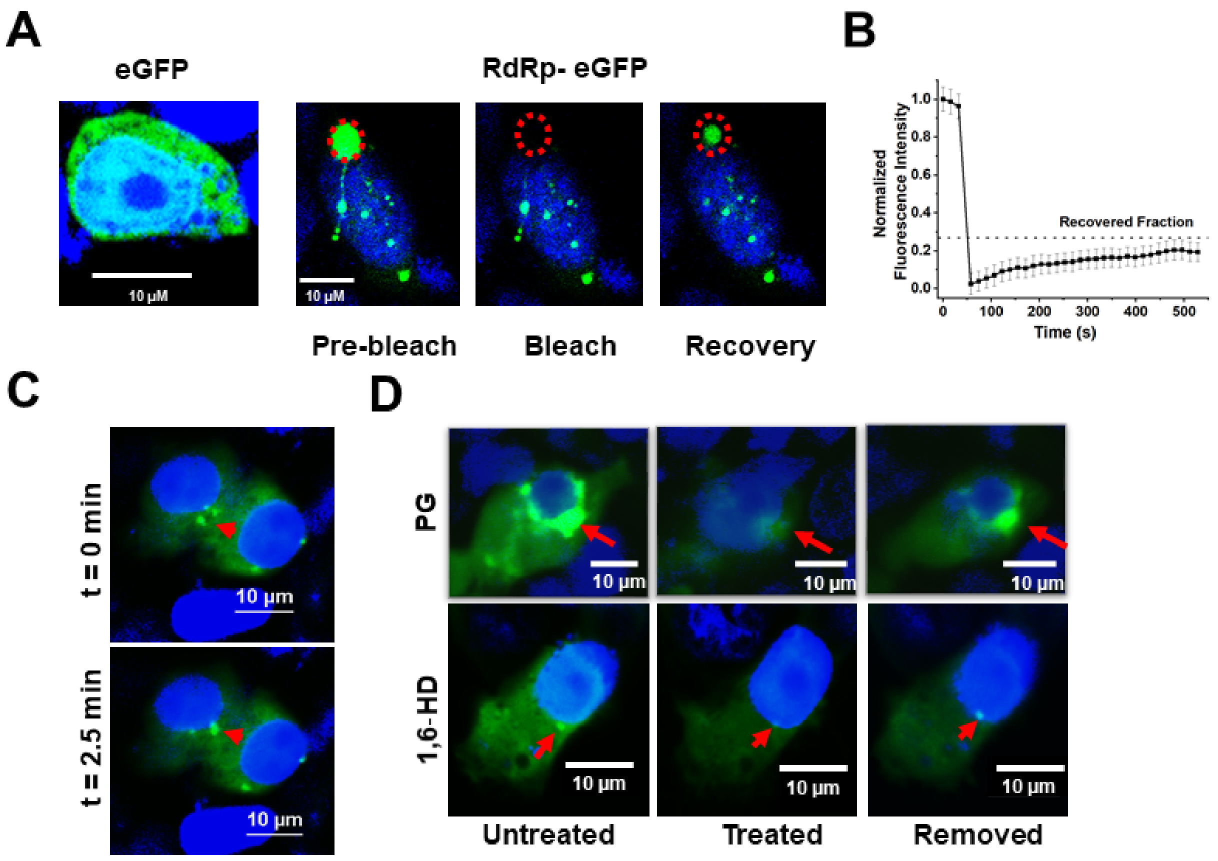


**Figure S9**. **RdRp forms LLPS in GII.4 transfected HEK293T cells. (A)** Transfection of RdRp-eGFP in HEK293T cells showing condensate formation. FRAP experiment was performed on one of the condensates (marked by a red circle), which shows the recovery of the after photobleaching. The eGFP-transfected HEK293T cells were used as a negative control where GFP is diffuse. **(B)** The FRAP graph shows the recovery after photobleaching. **(C)** Reversibility of dissolution of RdRp condensates with aliphatic alcohol 1,6 HD and PG. **(D)** Fusion of condensates in time-lapse imaging.

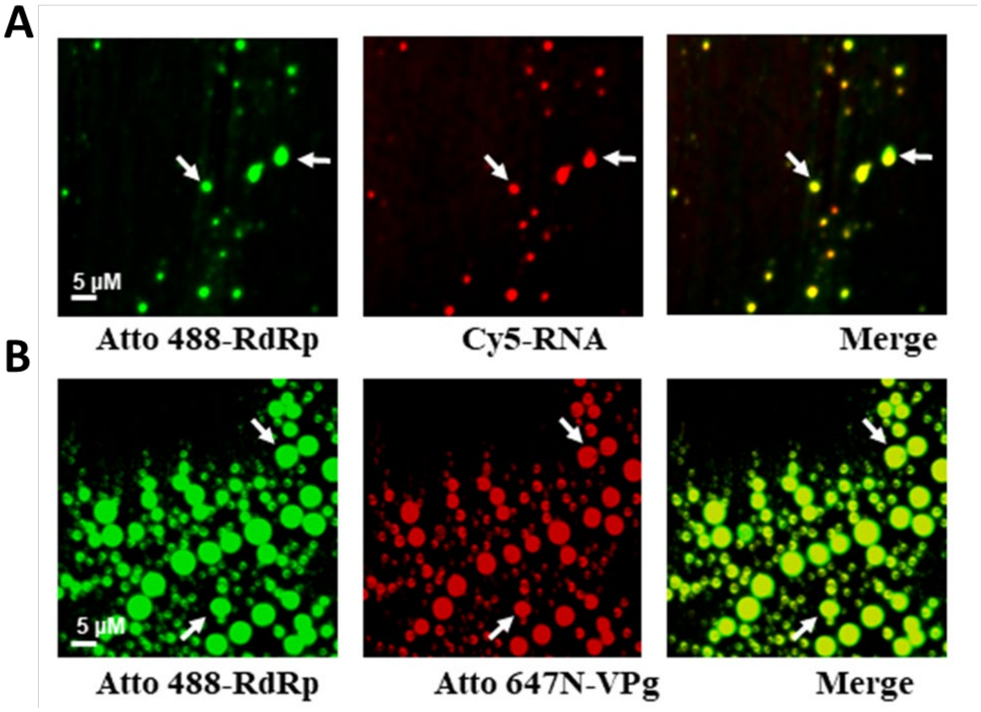


**Figure S10. GII.4 RdRp condensates sequester components of the viral replication machinery.** Confocal images showing the recruitment of **(A)** Cy5-RNA and **(B)** Atto 647N-VPg into the spherical condensates formed by Atto 488-RdRp. Colocalization of Cy5-RNA and Atto 647N-VPg are yellow in the Merge panel.

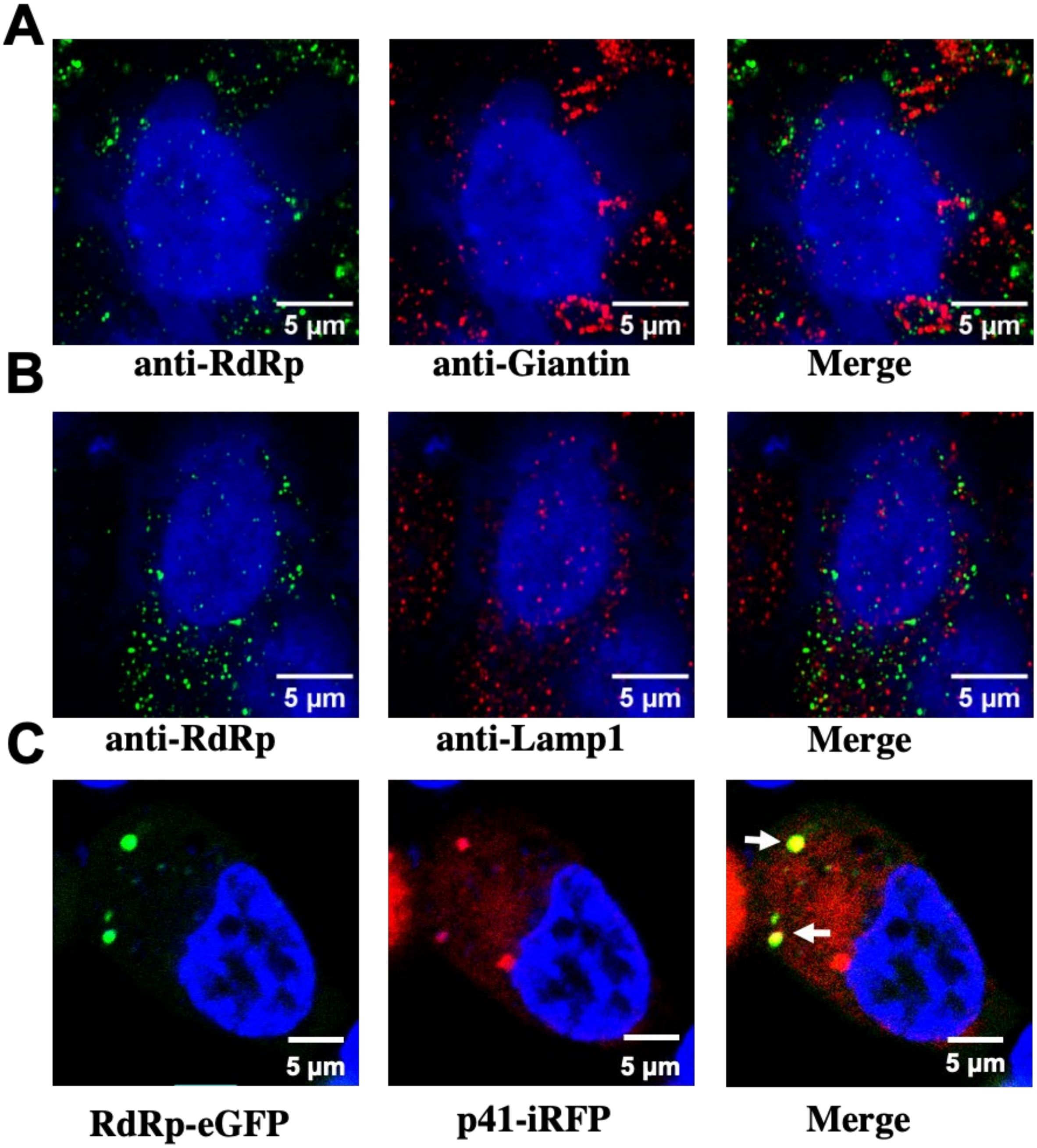


**Figure S11. Association of RdRp condensates with membrane markers.** GII.4 HuNoV-infected HIEs were fixed and dual labeled with markers for RdRp (green) and various cellular membrane markers (red) **(A)** *cis*/*medial*-Golgi (Giantin) and **(B)** lysosomes (LAMP1). **(C)** Colocalization (yellow) of p41-iRFP (red) and GII.4 RdRp-eGFP (green) was seen on overexpression of these proteins in HEK293T cells.

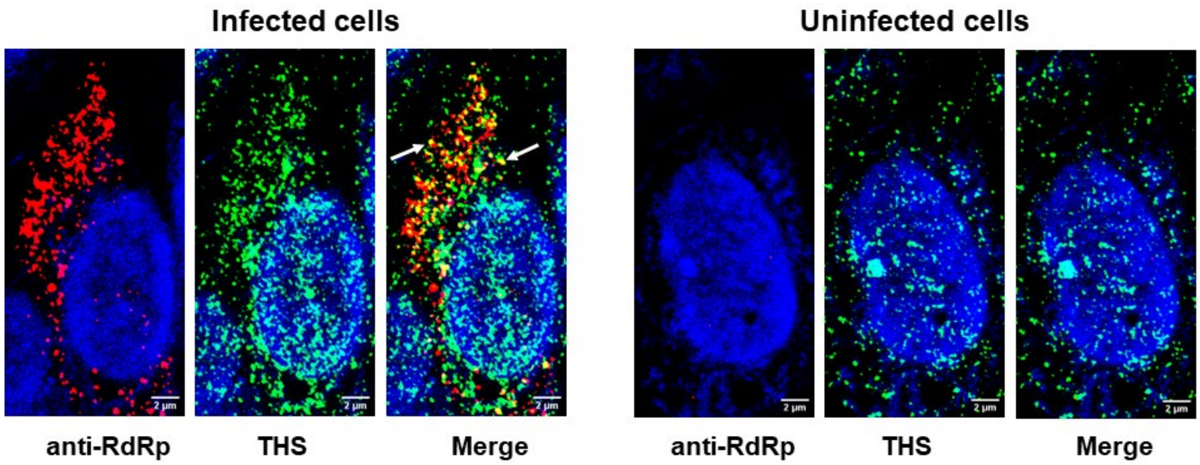


**Figure S12**. **Amyloid formation during virus-infection.** Left panel: Confocal microscopy images of GII.4 HuNoV infected HIEs labeled with anti-RdRp antibody (red) and stained with the Thioflavine S dye (THS, green). Right panel: Confocal microscopy images of uninfected cells.

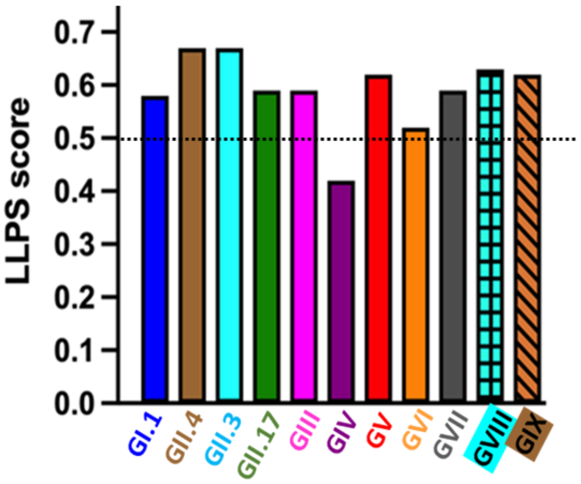


**Figure S13. DeePhase analysis shows RdRp sequences from different norovirus genogroups have the propensity to form LLPS**. The norovirus genogroups and their scores are indicated in different colors. The dotted black line indicates the threshold LLPS score of proteins to undergo LLPS.

### Supplementary video legends

**Supplementary Video 1: Dynamics of GII.4 RdRp molecules in condensates**

Video of the FRAP experiment showing the recovery of the RdRp molecules after photobleaching indicating the dynamic nature of these molecules in the condensate. In video, 1 sec = 30 sec

**Supplementary Video 2: Fusion of GII.4 RdRp condensates**

Time-lapse video of RdRp condensates showing the fusion of small condensates to form a bigger condensate. In video, 1 sec = 15 s

**Supplementary Table 1**: **Data collection and refinement statistics**

| | **GII.4 RdRp** |
|---|---|
| **Data collection** | |
| Space group | $P2_12_12_1$ |
| Cell dimensions | |
| *a*, *b*, *c* (Å) | 83.07, 83.70, 90.72 |
| α, β, γ (°) | 90, 90, 90 |
| Resolution (Å) | 37.21-2.03(2.10-2.03) * |
| Wavelength (Å) | 1.0 |
| $R_{sym}$ or $R_{merge}$ | 0.27(7.5) |
| $I / \sigma I$ | 8.4(0.81) |
| Completeness (%) | 99.86(99.27) |
| Redundancy | 11.4(10.5) |
| **Refinement** | |
| Resolution (Å) | 2.03 |
| No. reflections | 41484 |
| $R_{work} / R_{free}$ | 0.21/0.24 |
| No. atoms | |
| Protein | 3991 |
| Ligand/ion | 33 |
| Water | 158 |
| *B*-factors (Å$^2$) | |
| Protein | 44.09 |
| Ligand/ion | 56.32 |
| Water | 47.07 |
| R.M.S. deviations | |
| Bond lengths (Å) | 0.005 |
| Bond angles (°) | 0.67 |

*Values in parentheses are for highest-resolution shell.
